# Time-series sewage metagenomics can separate the seasonal, human-derived and environmental microbial communities, holding promise for source-attributed surveillance

**DOI:** 10.1101/2024.05.30.596588

**Authors:** Ágnes Becsei, Alessandro Fuschi, Saria Otani, Ravi Kant, Ilja Weinstein, Patricia Alba, József Stéger, Dávid Visontai, Christian Brinch, Miranda de Graaf, Claudia M E Schapendonk, Antonio Battisti, Alessandra De Cesare, Chiara Oliveri, Fulvia Troja, Tarja Sironen, Olli Vapalahti, Frédérique Pasquali, Krisztián Bányai, Magdolna Makó, Péter Pollner, Alessandra Merlotti, Marion Koopmans, Istvan Csabai, Daniel Remondini, Frank M. Aarestrup, Patrick Munk

## Abstract

Sewage metagenomics has risen to prominence in urban population surveillance of pathogens and antimicrobial resistance (AMR). Unknown species with similarity to known genomes cause database bias in reference-based metagenomics. To improve surveillance, we designed this study to recover sewage genomes and develop a quantification and correlation workflow for these genomes and AMR over time. We used longitudinal sewage sampling in seven treatment plants from five major European cities to explore the utility of catch-all sequencing of these population-level samples.

Using metagenomic assembly methods, we recovered 2,332 metagenome-assembled genomes (MAGs) from prokaryotic species, 1,334 of which were previously undescribed. These genomes account for ∼69% of sequenced DNA and provide insight into sewage microbial dynamics.

Rotterdam (Netherlands) and Copenhagen (Denmark) showed strong seasonal microbial community shifts, while Bologna, Rome, (Italy) and Budapest (Hungary) had occasional blooms of *Pseudomonas*-dominated communities, accounting for up to ∼95% of sample DNA. Seasonal shifts and blooms present challenges for effective sewage surveillance. We find that bacteria of known shared origin, like human gut microbiota, form communities, suggesting the potential for source-attributing novel species and their ARGs through network community analysis. This could significantly improve AMR tracking in urban environments.

## Introduction

Untreated sewage is increasingly becoming an important surveillance matrix for anonymized monitoring of large urban populations and has been used to monitor and quantify illegal drug consumption^1^, antimicrobial resistance (AMR)^2,3^, the bacteriome^4^, virome^5^, human population genetics^6^ and more recently SARS-CoV-2^7^. Different methodologies can be applied, but metagenomics has the advantage of simultaneously generating data on diverse genomic contents of a sample and thus provides an opportunity to simultaneously survey bacteria, parasites, viruses and AMR.

However, metagenomic data from untreated sewage are highly complex, reflecting a broad diversity of microbes shed by humans, as well as a high variability of sequences from plants, animals and unknown microbes, complicating the identification of relevant pathogen signals. The composition in the sewage is also influenced by different types of household waste (food), waste from individuals and external environmental sources (soil, groundwater) and microorganisms multiplying in the sewer system. Human fecal-associated bacteria account for 15-30%^8,9^ of the sewage influent community, meaning the majority of the sewage microbiome comes from other sources. Many bacteria in sewage are more permanent residents of the sewer system found in biofilms (*Pseudomonas, Lactococcus, Longilinea, Trichococcus, Acidovorax*)^10^, on the sewer pipe wall and in sediments at the bottom of the pipes (*Acrobacter, Acinetobacter, Aeromonas, Trichococcus*)^11^.

Starting in 2025, the European Union is looking to implement sewage-based surveillance in treatment plants serving 100,000 or more residents^12^. To perform reliable sewage surveillance AMR of human populations, it would be beneficial to first thoroughly identify the diversity in such cities, including their novel species, and composition changes through time. By doing so, we can establish a baseline of what is considered normal, which helps detect deviations that may indicate the presence or emergence of AMR. Identifying novel microbial species is also crucial as they may harbor unknown resistance genes, potentially revealing new sources of AMR. Since environmental bacteria inherently possess numerous AMR genes, distinguishing the origins of these genes is vital for the focused monitoring of AMR genes associated with human activity.

Previous studies have shown that the microbial composition within sewage can exhibit significant variability both temporally and spatially within the same sewage system^13^. In addition, there may be differences in composition and seasonality of microbes across different geographic regions^2^. These sources of variation occur on a time scale spanning from weeks to months. Finally, in order to compare sewage samples, many factors can be considered important, including temperature^9^, rainwater inputs^14^ and the size of the human population^8^ contributing to a specific sewage system community composition.

Traditional methods of studying microbes have relied on culturing them in laboratory settings and sequencing isolated cultures. However, this approach is limited because a significant portion of the bacteria in the environment cannot be cultured with traditional techniques^15^. Consequently, the genomic sequences of many taxa have not been discovered and are not represented in reference sequence databases^16^. Reference-based metagenomic approaches rely heavily on available references, posing challenges when analyzing samples containing a high proportion of uncharacterized taxa.

Yet, with adequate depth of the metagenomic sequencing, microbial genomes can be reconstructed directly from metagenomes^17^. Shotgun sequencing reads can be assembled to longer contigs and these can be grouped into genomes using shared characteristics such as co-occurrence and the frequency of short kmers. This process, known as genome binning, can yield draft or complete genomes, bringing to light previously unknown microbial species without the need for culture-based methods^17^. Without culturing, we use groups of 95%+ genome identity as proxies for species.

It is also important to consider that all metagenomic abundance data are compositional in nature because they capture the relative proportions of DNA fragments from various taxa instead of their absolute numbers^18^. This characteristic stems from the fact that modern shotgun sequencing operates with a fixed-capacity output, meaning the detected abundance of one taxon is dependent on and constrained by the collective abundance of all other taxa^19^.

With an increasing number of targets and epidemiological signals from mutations and flanking genes, metagenomics offers some interesting benefits compared to qPCR, but there are also difficulties. We here report on our efforts to retrieve and quantify thousands of bacterial species’ genomes from sewage and relate the results to the objective of AMR surveillance. We find huge spikes of environmental bacteria that challenge metagenomics surveillance only in some cities, but also find evidence that community detection of metagenomics data could aid the source attribution of species and AMR genes into human- and environmental-derived components. The sewage was derived from five major European cities over two years: Copenhagen, Denmark (Avedøre - RA, Damhusåen - RD, Lynetten - RL); Rotterdam, The Netherlands; Bologna, Italy; Rome, Italy; and Budapest, Hungary, encompassing seven different treatment plants (referred to as sites).

## Results

### Abundance and diversity of recovered genomes

Metagenomic assemblies of all the individual samples resulted in 42,731,574 contigs with an average length of 2,179 bp (N50: 2,303) and are in the following referred to as the single-sample assembly. Metagenomic co-assemblies of time points from the same site resulted in 25,120,037 contigs with an average length of 2,333 bp (N50: 2,569).

A total of 23,082 bins were recovered from kmer/depth-binning the individual metagenomic assemblies and 11,643 kmer/depth-co-abundance binning of the metagenomic assemblies. In total, 2,523 bins were found with completeness >90% and contamination below <5% and 12,687 bins with completeness >=50% and contamination <=10% according to CheckM2^20^.

We aligned ∼69% of the sequencing reads to the collection of species-level genome collection. When exploring the genomes’ abundances, we observed that in some samples, the genus *Pseudomonas_E* (GTDB-tk splits the wide *Pseudomonas* into new genus-sized genera) was highly abundant. This is primarily attributed to 12 different known *Pseudomonas_E* species (relative abundance > 0.05), including *Pseudomonas_E bubulae*, *Pseudomonas_E lundensis* and *Pseudomonas_E paraversuta*. None of our recovered *Pseudomonas* genomes belonged to the well-known opportunistic pathogen, *Pseudomonas aeruginosa*.

Alpha diversity (Shannon index) remained relatively consistent across Copenhagen and Rotterdam. However, notable alpha diversity drops were seen in sites with occasional *Pseudomonas_E* dominance (**Figure 1a**). See **Figure S1** for a more detailed timeline of *Pseudomonas_E* blooms.

**Figure 1.**
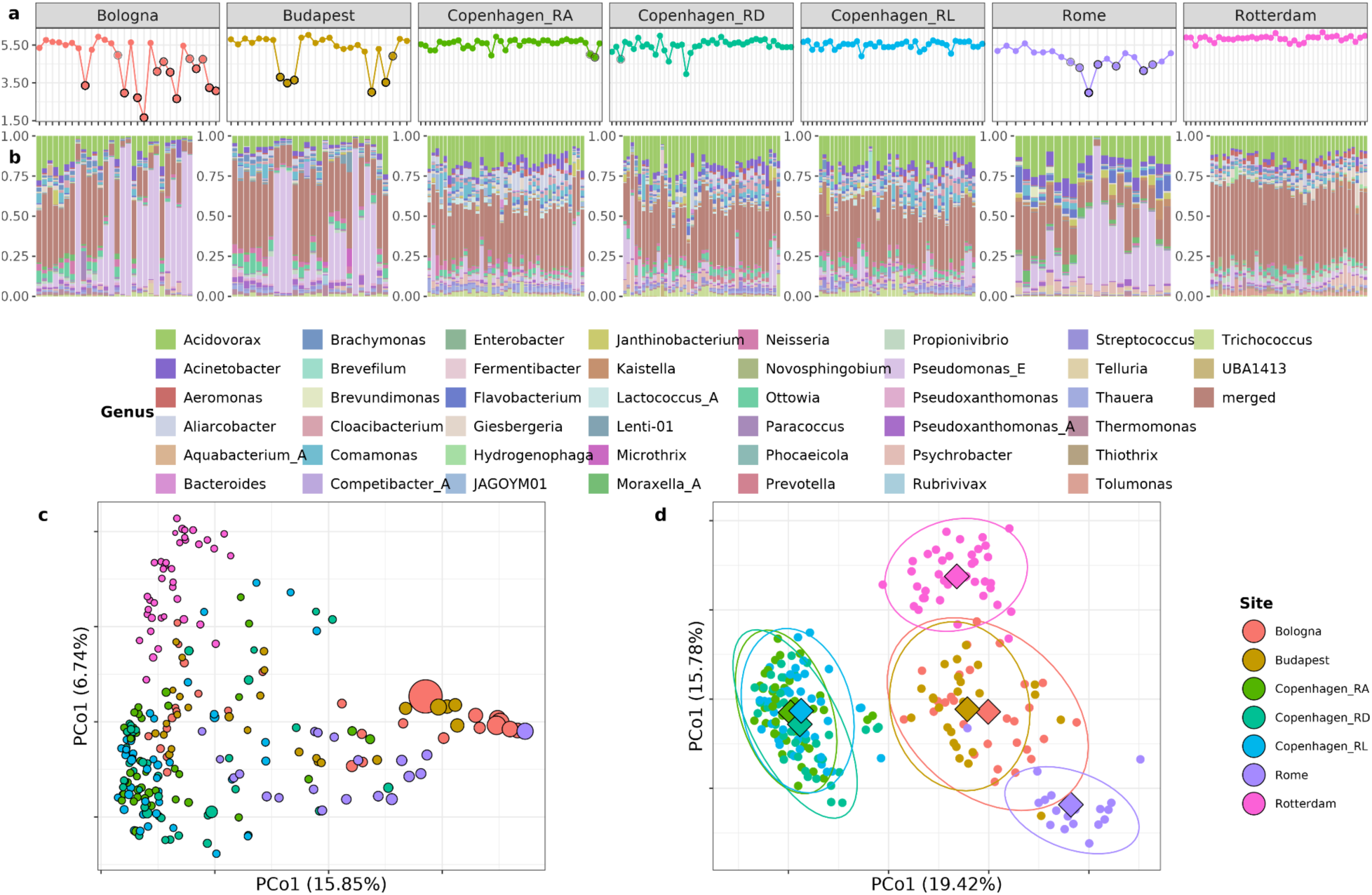
Abundance and diversity of recovered genomes through time and space. **a)** Shannon alpha diversity index of each sample. Circled dots indicate high levels of *Pseudomonas_E*. **b)** Stacked bar plots of the relative abundance of genera, stratified by sampling site and left-to-right sorted according to sampling date. The ‘merged’ category includes all remaining genera, for which no genus exceeded 5% of any sample. **c)** Multidimensional scaling of sample beta-diversity, based on Bray-Curtis (BC) dissimilarity. Point size is proportional to the inverse Shannon index. d) Also multidimensional scaling, but using the Aitchison distance instead of BC. Centroids are marked with a diamond and surrounded by an ellipse. These ellipses represent the 95% confidence intervals of the t-multivariate distribution for each sample group, serving as a graphical summary of the group’s variability.

The Bologna sewage microbiome exhibited notable changes over time, with certain genera appearing and disappearing through time. In Budapest the sewage microbiome was characterized by a consistent set of bacterial genera, however their abundances exhibited sample-to-sample fluctuations. In the Copenhagen sewage treatment plants, variations over time were more pronounced. For instance, the genus *Aliorcobacter* emerged in the second year of sampling in the RD and RL plants but appeared to be absent in the RA plant. In the last few samples from Rotterdam, the genera *Psychrobacter* and *Giesbergeria* became more abundant (**Figure 1b**).

In the beta diversity analysis using Bray-Curtis dissimilarity, the first principal coordinate predominantly captured the extensive diversity across the samples. Notably, it highlighted the samples primarily composed from the genus *Pseudomonas_E* associated with low diversity (**Figure 1c**).

The application of the Aitchison distance instead (see methods) highlighted the unique microbial signatures associated with each European city (**Figure 1d**). The three Copenhagen sites appear completely superimposed on each other, demonstrating low intra-city variability. To explore the variability of beta diversity, we found that the site explained ∼42% of the variance, while the city explained ∼38% (p ≤ 0.001). This suggests that the specific treatment plant location has a minor impact on the observed beta diversity, with the broader geographic location accounting for a more substantial portion of the variation, consistent with **Figure 1d**.

Furthermore, there is a significant overlap observed between the cities of Bologna and Budapest, highlighting a higher degree of similarity between these cities, than Bologna and its fellow italian city Rome.

### Co-assemblies especially boost recovery of rare and undiscovered sewage taxa

Utilizing a combination of co-assembly and single-sample assembly strategies we successfully retrieved 2,332 species of at least medium quality according to the MIMAG criteria suggested by Bowers et al.^21^. Many genomes lacked conserved RNAs but 56 were considered high-quality. MAGs recovered through single-sample assembly were generally more abundant (26% of the genomes exceeding 1% abundance), whereas only about 2% of MAGs recovered through co-assembly had at least 1% abundance in any sample.

Among the diverse orders represented in the genomic collection, Burkholderiales stands out with 410 different species. It is followed by the Pseudomonadales (148 species) and Bacteroidales (121 species) (Figure 2a). However, we were only able to assign species annotation to 988 of the species suggesting a substantial number of novel taxa.

**Figure 2.**
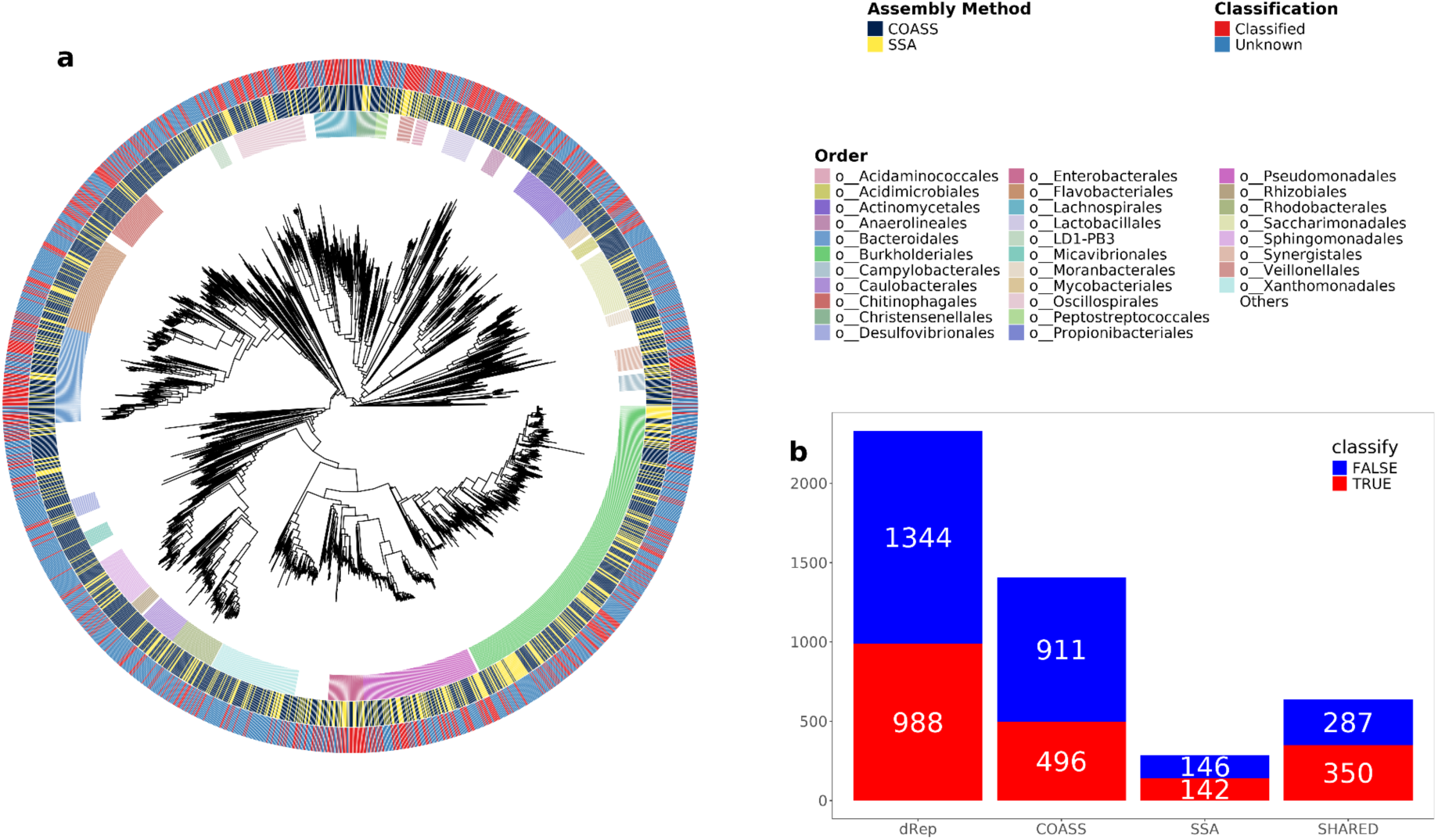
Thousands of species’ genomes recovered *de novo* from sewage. **a)** Phylogeny of all highest-scoring members of species-level MAG clusters. From inner to outer ring, they show genome-assigned taxonomic order, the assembly method employed and whether a genome was classifiable at species-level. **b)** A barchart showing the number of classified and unclassified species from one or both assembly methods. The SHARED category refers to MAGs that were recovered using both the single-sample assembly (SSA) and the co-assembly approach (COASS). Classify refers to whether GTDB-tk assigned species-level classification.

Using the single-sample assembly approach, we managed to recover genomes from 925 species (Figure 2b). The addition of co-assemblies significantly expanded the number of recovered species’ genomes with 1,407, to 2,332. Within this expansion, we identified 496 known species (Figure 2b). Notably, some well-known species, like *Lactobacillus acidophilus* appeared only through co-assembly. Nineteen species couldn’t be assigned to known orders, and intriguingly, 15 of those were exclusively obtained through the co-assembly process, indicating co-assembly can help recover substantially more novel taxonomy from sewage.

### Community detection in network graphs can assist source attributing genomes and AMR

We probed the co-occurrence of microbial taxa using network theory to identify their collective interactions. We analyzed the networks derived from each treatment plant separately, where each node (vertex) corresponds to a recovered genome and each edge (link) is a statistically significant co-occurrence of two species (Figure 3a).

**Figure 3.**
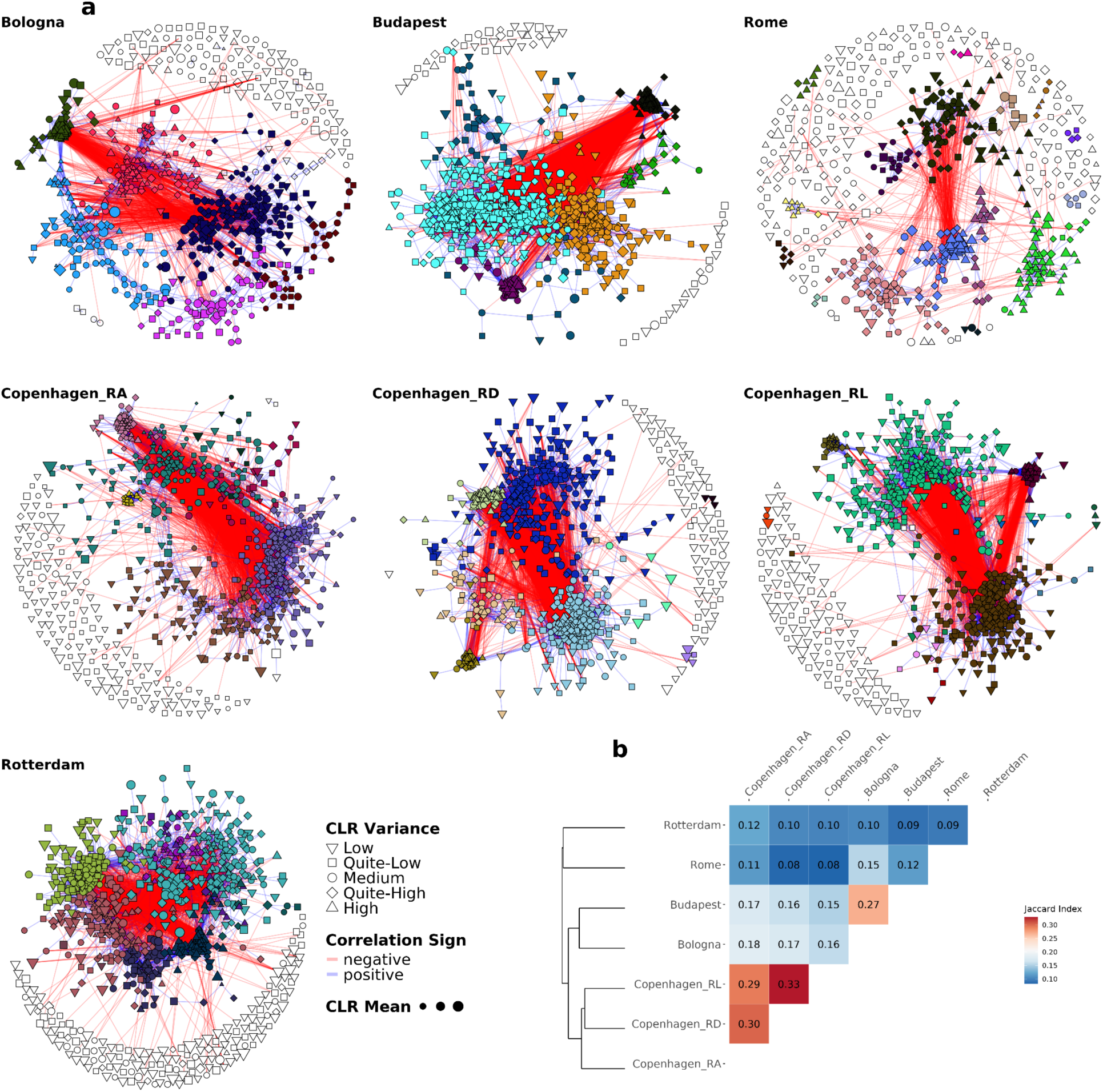
Diverse patterns emerge in the co-abundance networks of bacterial species across cities. **a)** Each vertex corresponds to a species’ genome, and the links represent significant positive (blue) and negative (red) correlations. The size of the vertices is proportional to the mean CLR-transformed depth of coverage, and the shapes encode variance level. Vertex colors are used to highlight the community membership. For the vertex placement, we used the Fruchterman-Reingold layout algorithm on the subgraph composed from only positive edges. Following this representation, many of the blue/positive links are covered by the circles, even if there are more positive links overall, see **Table S1**). **b)** Heatmap of the Jaccard similarity (J) with the hierarchical clustering of the Jaccard dissimilarity (1-J). The index J is defined as the ratio of the intersection of the links shared between two networks with respect to the union. See **Figure S4** for an analogous version with nodes colored by taxonomy.

Due to the absence of a ground truth and the numerous species under investigation, including a significant proportion of unclassified taxa, we opted to use straightforward correlations to construct the network. While this simplification may overlook finer details^22^, we specifically utilize correlation values deemed significant to assign weights for the links in our network analysis. This includes both positive and negative correlations, as detailed in the methods section, providing a quantitative description of the collective properties.

Our investigation unveiled striking disparities in network compositions and interactions, each manifesting a unique city-specific signature. The networks not only varied in the nodes/species constituents but also exhibited differing edge densities, as shown in **Table S1**. The number of distinct species remaining after filtering low-abundance measurements exhibited substantial variation, ranging from 546 species in Rome to 854 species in both Budapest and Rotterdam (**Figure S2**). Even more rare are the significant relationships common to all sites, with just 36 shared links. Notably, these shared connections consistently belong to a bacterial community, primarily composed of bacteria from the *Pseudomonas_E* genus **(Figure S3**). This *Pseudomonas_E* group, consistently present in all the sewage samples, is what we refer to as the PEH (*Pseudomonas_E* Heavy) community.

When we compare the sites in pairs, despite such variation, it becomes easier to capture common trends. Specifically, the three Copenhagen sites (RA, RD, and RL) displayed the highest degree of similarity among themselves, as substantiated by the Jaccard index computed on network edges (Figure 4b). Equally intriguing was the notable overlap observed between the network structures of Bologna and Budapest, while Rotterdam exhibited a distinct profile distinct from the other cities. These observations are consistent with the beta-diversity analysis (Figure 1c-d).

**Figure 4.**
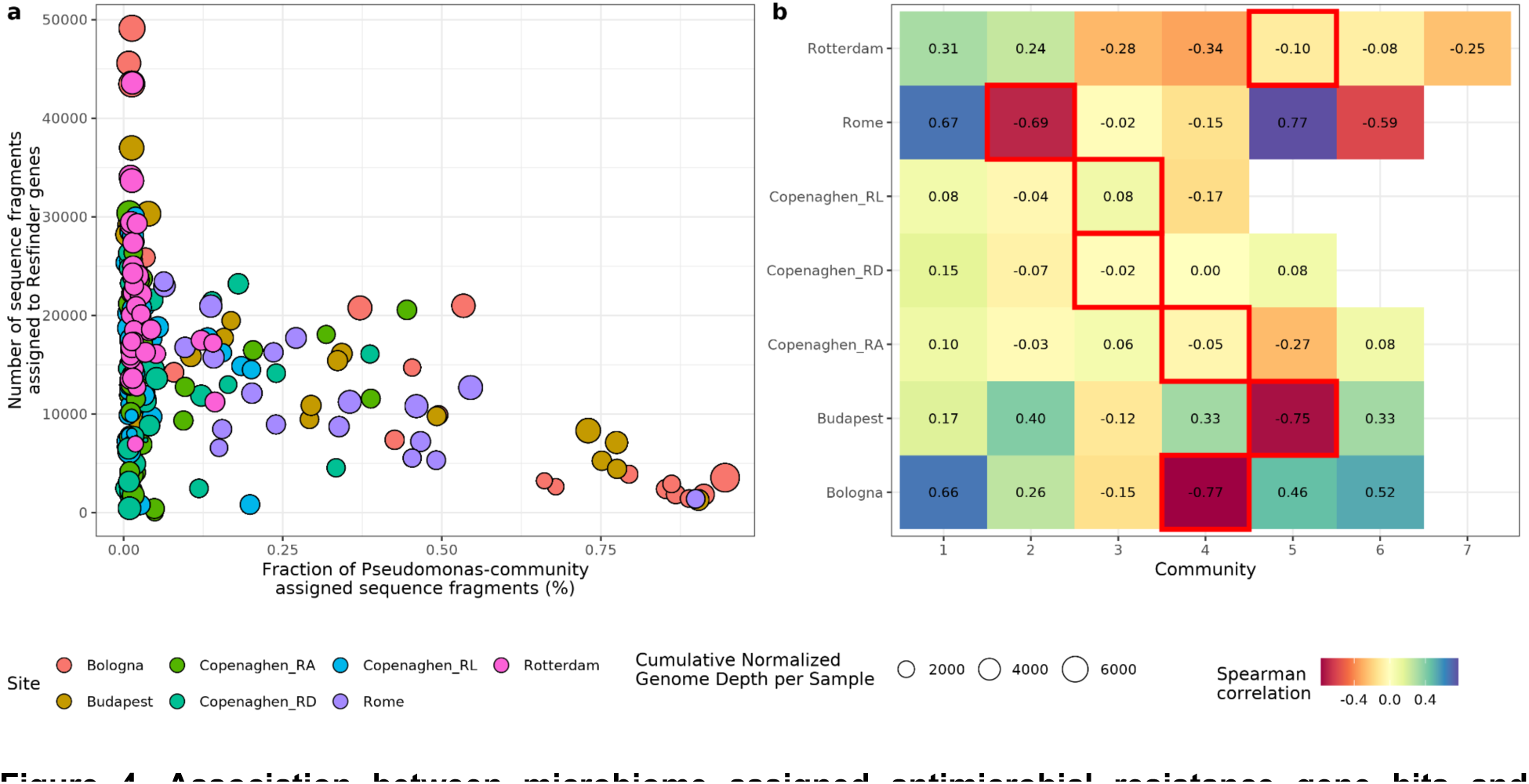
Association between microbiome assigned antimicrobial resistance gene hits and microbial community abundances. **a)** This scatter plot shows the inverse correlation between the fraction of sequence fragments attributed to *PEH* communities and the number of fragments assigned to ResFinder AMR in samples. Points are colored by site and sized according to the cumulative normalized genome depth per sample. **b)** Heatmap depicting Spearman’s ρ between the number of AMR-gene-attributed fragments and PEH community depth. Highlighted within red square rectangles are communities with heavy PEH species contribution.

We observed that networks consistently manifest a clear graph subdivision into distinct communities, as proved by their modularity values that always exceed 0.5 (**Table S1**). This network partition can be seen in Figure 4a, where bacteria cluster into discernible communities with strong internal connections rooted in only significant positive correlations.

Certain communities functioned independently with minimal interconnections. Others displayed numerous external connections, primarily negative, indicating competitive interactions. This enabled us to cluster bacteria into distinct communities and investigate their composition and temporal trends. Clear illustrations are the PEH communities observed in Bologna_4, Budapest_5, Rome_2, Copenhagen_RL_3, Copenhagen_RA_4, Copenhagen_RD_3, and Rotterdam_5, which are also the rare yet significant relationships shared across all sites (**Figure S5**).

Furthermore, examining the abundance trends of these communities reveals intriguing behaviors. Some communities exhibit clear oscillations synchronized with the solar year, as we will delve into in the next paragraph. Others display remarkable shifts in their relative abundance and dominance within the sewage composition, transitioning from low abundance to becoming predominant, as observed in communities like Rotterdam_4.

In the most extreme instances, PEH communities account for ∼95% of the assigned reads. We find a powerful, inverse relationship between the abundance of those communities and the observations of antimicrobial resistance genes (Figure 5a). The ratio between those two variables can vary by more than 1,000, suggesting environmental microbes can completely overpower surveillance targets in some cities.

**Figure 5.**
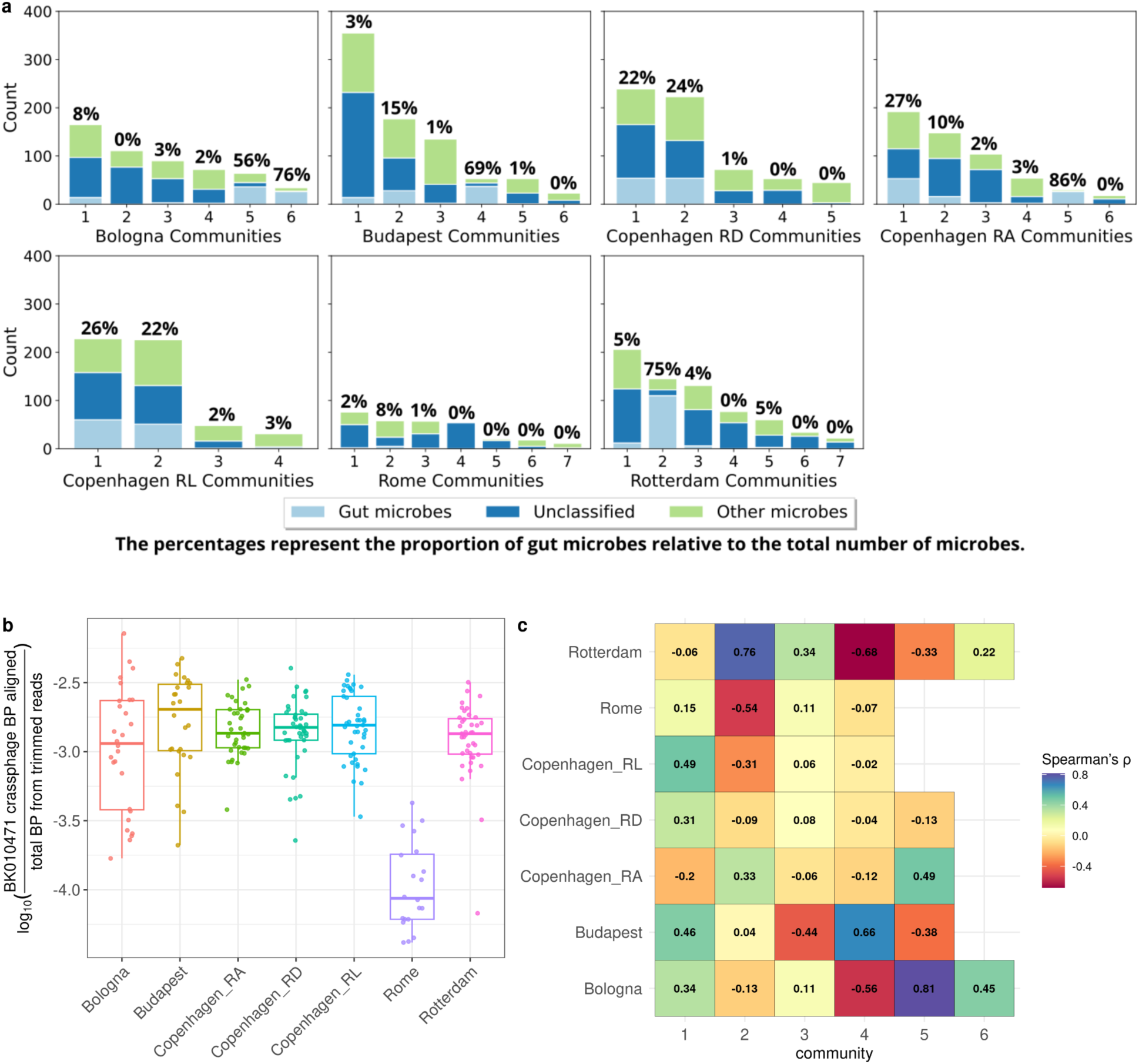
Human gut microbiome species form communities and correlate with crAssphage abundance. **a)** Certain sampling sites show communities dominated by gut microbes. Light blue indicates the number of MAGs classified as human gut microbes, dark blue represents the number of MAGs not identified at the species level, and light green represents the number of other species in each community. Communities with fewer than 10 members were excluded from the analysis. The percentage values indicate the proportion of human gut microbes within each community. **b)** Relative abundance of crAssphage (BK010471), an indicator of human fecal contamination shows low median level of fecal contamination in Rome but similarly high level of median abundance at other sites. **c)** Spearman’s ρ between sample crAssphage depth and CLR-transformed depths of bacterial communities.

The three Copenhagen PEH communities share most features, like having noticeable *Janthinobacterium tructae* participation absent from other communities (yellow). But they also show within-city differences, like the Copenhagen_RA community uniquely being associated with an unclassified *Giesbergeria* species (purple).

We correlated the AMR measures in samples with the abundance of bacterial communities and found it strongly anti-correlated specifically to the PEH communities in the three heavily affected cities (Figure 5b).

In an attempt to discover the drivers of *Pseudomonas_E* blooms, we associated the proportion of *Pseudomonas_E* to pH and temperature in the affected sites (**Figure S6**). There were no clear associations between those variables. The species-level beta-diversity was partly explained by temperature (8%, p=0.01) but not by pH (1.4%, p = 0.175).

#### Metabolic collaboration does not explain the observed sewage microbial communities

In an attempt to see if metabolic collaboration drove community formation, we investigated the degree to which the communities (or shuffled groups of the same size) encoded the proteome required for metabolic functionality.

Metabolism of most amino acids and central carbon metabolism, was common among all communities (including original and shuffled) and sites, while others, like plant pathogenicity or sterol biosynthesis, were completely absent. Sulfur biosynthesis, and nitrogen metabolism were more prevalent in some communities while absent in others, including shuffled ones, suggesting that these traits might be specific to a few bacteria rather than a collaborative effort among bacteria in the community.

As the community size grows, different metabolic categories tend to be more complete. However, there are some exceptions. For instance Copenhagen_RL_15, the smallest community in Copenhagen_RL with only two bacteria, still has complete histidine metabolism and sulfur metabolism. One of these bacteria, the *Thiotrix unzii* is well-documented for its involvement in sulfur metabolism^23^. In case of nitrogen metabolism, Rotterdam_7 is the only one which seems to have all nitrogen metabolism modules.This community consists of 22 bacteria most of them are not classified at the species-level. Among them, one MAG is classified as the genus *Nitrospira_F,* which is known for its nitrification capabilities^24^. For a breakdown of community metabolic differentiation, see **Figure S7.**

#### Human gut microbiota cluster into fecal communities

Yet another explanation could come from the fact that sewage is a mixed environment with multiple input sources. When these vary in relative contribution, all organisms from the same source will co-occur also leading to groups of genomes co-occurring through time. In essence, our communities would then be multi-genome bins, exploiting differential abundance in the same way as genome binning tools can, but with weaker associations.

If e.g. the amount of environmental bacterial contribution varies significantly through time, we would expect those genomes to group together and differently from human microbes.

By comparing our classified species to known human gut microbiota species. Out of 998 MAGs classified at the species-level 241 potentially originated from the human gut. We observed that these MAGs tend to form communities, Figure 5a. We noted that certain sampling sites exhibited communities dominated by gut microbes, such as communities 5 and 6 in Bologna, community 4 in Budapest, community 5 at Copenhagen RA, and community 2 in Rotterdam. Interestingly, in Rome, gut microbes are rare in the communities. Most sites have a single community with more than ⅔ MAGs being recognized as human fecal microbiota.

To further confirm this finding, we used crAssphage as an indicator for human fecal contamination (see Methods). With the exception of Rome, which exhibited low levels (corresponding with a lower count of human gut microbiota species within the communities), all cities showed similarly high median abundances of crAssphage, Figure 5b . By quantifying its abundance, a pronounced correlation emerged between crAssphage levels and the dominance of fecal bacteria within communities, Figure 5c. This is particularly evident in Bologna communities 5 and 6 (with Spearman’s correlation of 0.81 and 0.45), community 5 in Budapest (0.66), community 5 in Copenhagen RA (0.49), and community 2 in Rotterdam (0.76).

Additionally, we quantified human mitochondria and used those numbers to confirm that crAssphage abundance was highly correlated with human-associated DNA (**Figure S8**). Since crAssphage is an established fecal indicator with more aligned reads than mitochondria, it was expected to be associated with less stochastic sampling noise, and we chose to use it further in our analyses.

#### Community detection for assigning AMR to sources

Looking at Figure 4b, we saw several communities that are strongly positively correlated with AMR. However, these were not necessarily the fecal communities. We therefore sought to include the ARGs in the per-site network analysis and recomputed the networks with both ARG and MAG features included and correlated. While correlation alone might be too uncertain to ascertain source genome, an overall origin based on community might be feasible.

Using such an approach, Bologna, Budapest and Rotterdam formed a fecal community each with ARGs and at least 70% of classified species being known human gut species.

The Rotterdam fecal community was associated with 10 ARGs, mostly *tet*(W)/(O)/(32) alleles which are known to recombine, *aph*(3’)-III, *bla*ACI and *cfr*(C). The Bologna fecal community contained 18 genes with many recognized Proteobacteria ARGs like sul(1), *sul*(2), *qnr*S, *tet*(X), *tet*(W) and *aad*A alleles. Lastly, the Budapest fecal community included 22 ARGs, overlapping with some of the others, but also with unique inclusions like *mph*(N) and *tet*(44) and *erm*(B).

In the Copenhagen sites, there was a tendency of large ARG-only communities forming, not associated with any species in particular (**Figure S9**). Whether this is due to plasmid carriage or other sources remains to be explored.

### Some bacterial communities closely follow the solar year

To better visualize and understand the dynamics between the communities, we explored the temporal trends of microbial communities across multiple sites (Figure 6). As anticipated, the network structures unveiled complex interrelationships among these communities, reflecting significant changes in sewage composition over time. Notably, we identified communities characterized by strong periodic trends that exhibited clear negative correlations with other communities. In contrast, certain communities experienced a notable longer-term abundance change, going from the least to the most abundant community in a >1 year sampling period, or reversely, changing from dominant to insignificant.

**Figure 6.**
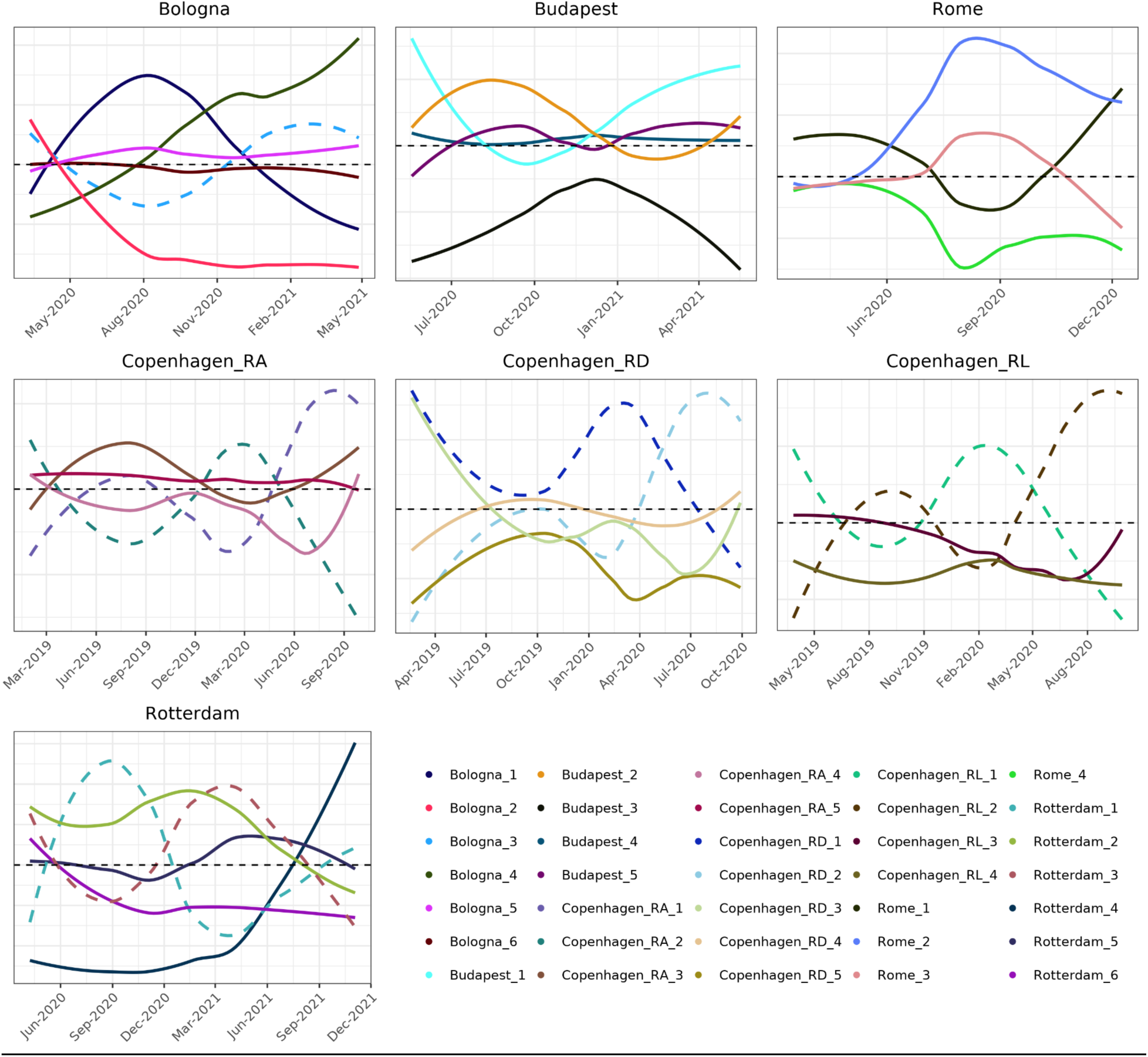
Time abundance of principal bacterial communities at each site. The figure shows the smoothed (LOESS) trends of the CLR-transformed depths of bacterial communities, colored as in Figure 3. The horizontal black dashed lines indicate where CLR-transformed depths are zero, highlighting the geometric means of the samples serving as the reference in CLR space. Only the communities composed of 30 or more bacterial species/MAG are included in the figures. Communities displaying periodic behavior aligned with the solar year are marked with dashed lines.

We quantified the time periods of the ’periodic’ bacterial communities. Given that our sampling campaign lengths frequently covered only a single oscillation per community, we opted to simply estimate the frequency by fitting sine functions. Notably, we identified a subset of communities, especially in Copenhagen, Bologna, and Rotterdam, which displayed periodic behaviors closely aligned with the solar year (**Figure S10**). The community most closely following the solar year was Copenhagen_RA_2 (364,77 +/- 9.48 days). This community contains *Acidovorax defluvii*, *Trichococcus flocculiformis* and 145 other species, most of which have not been described before.

Intriguingly, in Rotterdam and across the three Copenhagen sites, we discovered two annual communities that were markedly negatively correlated with each other. These are visually represented by dashed lines in Figure 6. Additionally, **Figure S11** provides another perspective by showing the trends in beta diversity per site over time. We observed a notable divergence in the bacterial composition of sewage at sampling sites over time, with samples becoming increasingly dissimilar, following time between collection dates (Δ days) (**Figure S11b).** Budapest, though, emerged as an exception as its sewage composition exhibited relative stability over the course of a year. In half of the sampling sites, there is a detectable decrease in beta diversity after one year. This particular phenomenon was absent in Bologna and in Rome. (**Figure S11a)**. Excluding the dominating *Pseudomonas_E* taxon made little difference, demonstrating that the seasonal findings are separate (**Figure S11c**). Although some sampling sites exhibited periodic patterns aligned with the solar year, others did not; the diagrams at the community level reveal more nuanced patterns within each site, distinguishing between communities with seasonal behaviors from those without.

## Discussion

We managed to uncover a large number of novel taxa, representing a wide phylogenetic span, from the sewage samples in five European cities. Many novel taxa, including orders, we could only recover by the combination of deep co-assemblies and co-abundance genome binning. It would seem that using single-sample pipelines only recovers a relatively small fraction of abundant taxa in sewage, and other tactics are required to discover the majority of new life. Downsides of single-sample binning were also highlighted in a recent investigation that identified large degrees of contamination of public MAGs generated with such approaches^25^. The large added depths of co-assemblies resulted in many contigs that had too low depth of coverage in the individual samples. Only ∼2% of species we recovered with co-assembly had at least 1% abundance in any of the samples, while species recovered with single-sample assembly were more abundant in general, highlighting the added sensitivity. A downsize was the extensive compute requirements with some jobs running for many weeks and needing a lot of memory. The many new taxa recovered here by reference–free metagenomics improve the public collections and will potentiate future reference–based metagenomics and sewage monitoring.

We found that sewage microbiomes drift apart over time so samples that are temporally close, also have lower beta-diversity confirming expectations and previous findings^13,26^. Surprisingly though, we found striking differences in the impact of seasonality on the surveyed cities. Some cities like Rotterdam and Copenhagen showed large effects with samples taken a year apart being much more similar. Other cities, like Bologna, however had almost no seasonal effects, with a more stable microbiome throughout the year. This result has great implications for current and future sewage surveillance and how those campaigns are designed. If we want passive low-frequency surveillance of e.g. AMR in sewage, like previously proposed, we need to ensure this is taken into account^2,3^. Perhaps the exact timing of sampling for a health snapshot is unimportant in one city, whereas it is paramount in another. As sewage surveillance of pathogens and AMR is increasingly used and soon mandated in the EU, these findings are important to take into account, especially if using a metagenomic approach.^12^

Network analyses reveal a separation of bacteria in distinct communities, that correspond to e.g. different sources of origin or seasonality / temperature. Cities with distinct seasonality exhibited large bacterial communities interconnected by negative correlations. These intercommunity relationships display strong seasonal fluctuations, which likely contribute to the observed beta-diversity. Also interesting is the existence of bacterial communities that drastically change and consistently increase or decrease over the entire sampling periods. This was particularly pronounced in Bologna and Rotterdam, where initially marginal communities assumed dominance, and vice versa. This underscores the fluid nature of microbial community dynamics. Our findings allude to a multifaceted microbial ecology in urban sewage systems, where community dynamics are shaped by a multitude of factors.

Furthermore, the finding that a great abundance of polymicrobial *Pseudomonas_E*-rich communities occasionally dominate sewage samples (>90% reads) should inform future surveillance efforts. Although a high abundance of *Pseudomonas* is not unprecedented in the literature^27^, the noteworthy occurrence of extreme blooming, quick fluctuations and multispecies nature stands out.

Potential factors driving these blooms include rainwater inputs, temperature and pH fluctuations, occasional nutritional starvation, and the role of these species in both the formation and detachment of biofilms. We found no connection between temperature or pH fluctuations and the occurrence of these blooms. As many *Pseudomonas* species frequently engage in biofilm formation, one might suspect that PEH corresponds to biofilm lining surfaces in the sewage system. Physical connectedness in a biofilm matrix would also appear to explain why specifically these communities are much more close knit (high co-occurrence) compared to other detected communities. If so, this type of analysis could be used to profile individual sources.

It would appear that one could source-attribute many novel species through network analyses and inferring physical proximity / multicellular behavior. Indeed, we also see that known human gut microbiota, human mitochondria and crAssphage correlate. If further refined, one could imagine a system where simply point sampling at the treatment plant through time can provide the data to differentiate changes occurring inside the human guts, rat guts, sewer biofilms and other input sources. While further examination of these factors should be a future focus, the domination of *Pseudomonas_E* species also has important implications in a surveillance context.

First of all, compositional data analysis (CODA) will be essential in any surveillance efforts to avoid spurious correlations. As previously demonstrated by Gloor et al., microbiomes are compositional, and the issues of non-CODA approaches become particularly obvious when a few actors, like our *Pseudomonas_E* species, dominate the samples.^19^ The foundation of CODA lies in considering the ratio between features, rather than their absolute values. Considering the sample dominated by *Pseudomonas_E* communities, the relative abundances of other components are compressed to increasingly lower values, making it challenging to discern their specific characteristics. However, when focusing exclusively on the ratio between bacterial species, they remain unaffected by this phenomenon. This is highlighted by the differences in Figure 1c and Figure 1d, where the former is primarily driven by *Pseudomonas_E*, thus concealing the city–specific species composition shown in the latter. Another approach attempting to circumvent the compositionality of microbiomes comes from absolute quantification, where each read-quantified entity is related to some additional measure like spike-in sequences or bacterial cells^28^.

Neither CODA, nor attempts to estimate absolute quantities from sequencing results can however solve *Pseudomonas_E*–related problems. As almost all recovered DNA suddenly belongs to an environmental *Pesuedomonas_E*-rich community, we will get very little data on pathogens and AMR which are surveillance targets of larger interest. Ratios between this community and resistance gene hits can in fact vary more than 1000–fold. Fewer discrete read counts to surveillance targets of interest mean larger stochastic sampling noise, lower sensitivity and more expensive surveillance. If *Pseudomonas* communities are from the biofilms and tend to shed in specific daily time slots, it might be worthwhile to target smaller time windows than that of 24-h continuous samplers. Prescreening samples for *Pseudomonas* with cheap qPCR or culturing could also help inform which samples should be subject to shotgun metagenomics.

Even though we surveyed a large number of samples across five countries, we only included European cities in this study. We know from previous work that historically undersampled geographic regions, from a genomic point of view, like Africa and South America, have a much larger proportion of metagenomic dark matter. Enteric pathogens and AMR are likewise problems that will impact the Southern Hemisphere to a higher degree in the future^29^. Whether our results are generalizable to all of Europe or the World should receive future attention.

In conclusion, there are still thousands of undiscovered microbial species in European urban sewage, a system that has recently undergone extensive studies. The use of more frequent shallower sequencing with co-assembly and co-abundance binning can help us uncover much of this diversity. The microbial communities frequently get completely overwhelmed by large multi-species, *Pseudomonas_E*-dominated blooms which need to be accounted for in untargeted surveillance systems on sewage DNA. Finally, we suggest that the structured microbial time-series communities we observe open up for possibilities that we can actually identify the physical source of novel species in e.g. biofilms and other microbiomes.

## Methods

### Sample collection

Sewage samples were collected as part of a longitudinal study across European cities (**Table S1**) with three sites in Copenhagen (Denmark), being Rensningsanlæg Avedøre (RA), Rensningsanlæg Damhusåen (RD) and Rensningsanlæg Lynetten (RL). Sites in Budapest (Hungary), Rotterdam (Netherlands), Bologna (Italy), and Rome (Italy) were also included. We sampled Copenhagen from 2019, and the other cities starting in 2020 (**Figure S12**)^30^. Each participating partner was responsible for collecting and shipping 1-2 liters of untreated urban sewage samples to Technical University of Denmark (DTU). Collection was performed using 24-hour automatic continuous-flow samplers used everywhere except in Bologna. In Bologna, three individual grab samples of 300 ml were collected manually 5 minutes apart in the 8 - 10 AM interval and pooled.

We used a weekly sampling frequency, but some time points were missed due to the COVID-19 lockdown. Sewage samples, collected and frozen at −80°C, were transported to DTU under IATA regulation SP A197 as UN3082, adhering to safety standards with volumes not exceeding 2L. At DTU, each sample received a unique identifier, linking it to essential metadata including location and collection date to ensure traceability. This process ensured compliance with the Danish Act on the scientific ethical treatment of health research, indicating no requirement for prior authorization from ethical review boards.

### DNA extraction, quantification and sequencing

500 mL from each sewage sample was defrosted over 2 days at 4 °C. Once thawed, the samples were centrifuged for 10 minutes at 10,000 x g, allowing for the collection of sewage pellets. These pellets, designated for DNA extraction, were stored at −80 °C.

Following the protocol established by Knudsen^31^ et al. and used later e.g. by Hendriksen et al.^2^., DNA was extracted from the sewage pellets. This validated approach for untreated sewage ensures the consistent and reliable recovery of metagenomic DNA with high microbial diversity. The extracted DNA was quantified using the Qubit 2.0 DNA HS Assay (Thermo Fisher Scientific, Waltham, MA), for assessing the quality and concentration of the genetic material obtained. Extracted and quantified DNA was shipped to Admera Health (New Jersey, USA).

At Admera Health, library preparation for metagenomic analysis was conducted using the KAPA Hyper PCR-free library preparation kit (Kapa Biosystems, Roche, Basel, Switzerland), following the manufacturer’s guidelines. The quality and quantity of the prepared libraries were then assessed using both the Qubit 2.0 DNA HS Assay and QuantStudio® 5 (Applied Biosystems, Foster City, CA), ensuring they met the standards required for Illumina sequencing. Sequencing was then performed on the Illumina NovaSeq6000 platform, targeting at least 35 million clusters with 2 x 150 cycles (paired-end reads), corresponding to 10.5 Gbp per sample.

### Genome recovery

Metagenomic reads were preprocessed by trimming sequences (Phred score < 20; minimum length of 50 bp) and removing adapters using bbduk2, which is part of the bbtools^32^ suite (v 36.49).

MEGAHIT^33^ (v1.2.8) with the “meta-large” preset and a minimum contig size of 1,000 was used to co-assemble the paired-end metagenomes in groups stratified by sites. In addition, to the co-assembly, Lazypipe^34^ (v2.1) was run on each of the sample sequence runs individually to acquire single-sample assemblies. Lazypipe is a virus-focused pipeline but initially runs the MEGAHIT metagenomic assembler (v1.2.9 with default parameters) as part of its workflow^34^.

Minimap2^35^ (v2.24) was used to align reads back to both the single-sample assemblies and co-assemblies, so that all sequence runs used to produce the initial assembly were also used for alignment. The resulting BAM files were processed with jgi_summarize_bam_contig_depths to generate single-library depth files for the single-sample assemblies and multi-library depth files for the co-assemblies.

MetaBAT2^36^ (v2.15) was then used to produce bins from each assembly, using its corresponding contigs and depth file. Only genome bins of at least 200,000 bp were retained as is the default and is enough for almost all bacterial genomes.

We aimed to construct a comprehensive, non-redundant, and environmentally representative reference genome dataset covering all the sewage samples. CheckM2^20^ (v1.0.1) was used on all 34,725 MetaBAT bins, originating from the two approaches: 23,082 single-sample assembly bins, and 11,643 co-assembly bins. Based on CheckM2 results, we retained genome bins with contamination ≤10% and completeness ≥50%, corresponding to the MIMAG Medium quality threshold^21^. 12,687 genomes fulfilled the criteria and were compared with the dRep^37^ (v3.4.2) *dereplicate* workflow to find the highest-scoring representative of each species cluster. Briefly, the dRep dereplication process first clustered the genomes based on their MASH^38^ distances, with a threshold set at 0.9 and a sketch size of 1000 and subsequently did secondary clustering with an average nucleotide identity (ANI) of 0.95. The process yielded 2,332 representative species-level MAGs.

GTDB-tk^39^ (v2.4.0 with GTDB r220 as reference) was used to classify the dereplicated MAGs and place them into the and archaeal phylogenetic trees, using the default values of min_perc_ac=10 (percent amino acids covered by a MAG) and min_af=0.5 (proportion genome coverage for species assignment).

For a visual summary of the main steps involved in this pipeline, see the flowchart in **Figure S13** in the Supplementary Information.

### Species quantification

All the trimmed, individual sequence runs were aligned to the dereplicated genome collection using minimap2^35^ with paired-end alignment, N=3 and otherwise default settings. Only the paired-end reads with both reads successfully passing the preprocessing step were aligned. The resulting BAM files were processed with the jgi_summarize_bam_contig_depths (v2) script and supplied to MetaBAT2 with default settings. We then multiplied the contig length out of each depth value to obtain the number of nucleotides aligned to contigs that could then be summed to its parent MAG in each sample. The depth was then re-computed for each sample-MAG combination.

We employed centered log-ratio (CLR) transformed abundances to handle the inherent compositional nature of the data^40^. To address cases where CLR is undefined due to zeros, we used a pseudo-count equal to 65% of the sample detection limit as previously recommended^41,42^.

### Beta diversity and metagenome ordination

We performed Principal Coordinates Analysis (PCoA) by utilizing the *`cmdscale`* function from the *`stats`* package in R^43^. PCoA is a dimensionality reduction technique that transforms a matrix of dissimilarities into a new coordinate space where axes, termed principal coordinates, represent the most substantial variance in the data. This transformation facilitates the visualization and subsequent analysis of complex datasets in lower-dimensional scatter plots.

To determine the structure embedded without the complex microbiomes, we applied two distinct dissimilarity measures. The first measure employed was the Bray-Curtis dissimilarity, calculated using the vegan (v2.6) package^44^. This metric is frequently used in quantifying ecological variation across samples, making it useful for identifying general patterns of diversity and distribution.

However, to address the unique challenges presented by compositional data, such as those inherent in metagenomic datasets, we also utilized the Aitchison distance. This metric is specifically tailored for compositional data analysis. By applying it to CLR-transformed data, as outlined in the preceding section, we could effectively capture the nuances of compositional variability. The Aitchison distance, fundamentally a Euclidean distance computed on CLR-transformed data, can uncover subtle yet critical compositional disparities between samples^45^.

To determine if sampling site and city have significant effect on beta diversity we performed a permutational multivariate analysis of variance (PERMANOVA) using the adonis2 function from the vegan^44^ (v2.6) package. This analysis was conducted on Aitchison distances.

### Network community reconstruction and analyses

Initially, we filtered out genomes with a mean depth of coverage of 0.5 or lower. Genome depths in each sample were CLR-transformed. Genome abundances were then correlated to each other using Pearson’s correlation and we retained genome-genome links with p-values =< 0.1 after correcting for the false discovery rate.

We defined our networks as undirected, weighted, and signed graphs, where genomes are vertices, and significant correlations are links. To extract communities we used a signed version of the modularity able to manage also the negative weights associated with the links^46^. Therefore, a community is the subset of bacteria densely connected from only positive correlation, minimizing the negative ones.

All network analyses were conducted using R^43,47^ through the package *mgnet*, available at https://github.com/Fuschi/mgnet, with key dependencies being the igraph^48^, ggraph^49^, qualpalr^50^ and psych^51^ packages.

### Metabolic potential of communities

The metabolic potential of distinct communities was measured using METABOLIC-G (v4.0) with the default options for MAGs and aggregated to community level. Additionally, we analyzed shuffled communities, where the same number of bacteria from the same site was randomly shuffled and reorganized into new communities. From the resulting table we utilized ‘KEGGModuleHit’ data. KEGG (Kyoto Encyclopedia of Genes and Genomes) modules were grouped into categories, and the number of modules present in each community per category was calculated. Subsequently, we determined the ratio of modules present out of the total modules in each category. KEGG module categories with at least one community with 0 completeness and KEGG modules with at least one community with minimum 0.7 completeness were visualized in **Figure S7.**

### Antimicrobial resistance gene quantification

Trimmed reads of all libraries were mapped using kma^52^ (v1.2.8) with the -cge and -1t1 flags using both paired-end and singleton reads as input against the ResFinder database^53^ (commit=3eedbde). The ResFinder database consists of a manually curated collection of antimicrobial resistance sequences^54^. Settings of kma allowed mapping only one query sequence per template and with default penalty values.

The aligned sequence fragments were summed to representative sequences in groups of similar ResFinder alleles. This clustering was accomplished using 90% identity and 90% coverage thresholds with usearch (v11.0.667) cluster_fast on ResFinder genes^55^.

### Associations to environmental and human fecal indicators

To identify the overlap of human microbiota and our classified species, we downloaded the genome collection from Unified Human Gastrointestinal Genome (UHGG) collection^56^ (v2.0.2) representing species derived from the human gut. We then determined the number of those species in our classified communities by their clustered name alone.

To calculate the abundance of the human mitochondrial genome, trimmed reads were aligned by minimap2^35^ (v2.24) to the human mitochondrial sequence (hs_ref_GRCh38.p7). The average alignment depth per sample was determined using the jgi_summerize_contig_depth function from MetaBAT2^36^ (v2.15).

To calculate the relative abundance of *Pseudomonas_E* in each sample, we summed the mean depth of each MAG within the *Pseudomonas_E* genus, and then divided this sum by the total mean depth of all MAGs.

The reference genome of *Carjivirus communis* BK010471 was downloaded from the NCBI nucleotide database using the Entrez Direct utility. Paired-end reads were aligned to this reference genome using BBMap, with a minimum identity threshold of 90%. Depth of coverage of the aligned reads was retrieved using Samtools^57^ (v1.19.2).

### Periodicity Analysis of Bacterial Communities

For periodicity analysis in urban sewage bacterial communities, we applied a sine wave fitting method post-CLR transformation. Our model, f(t)=A*sin(ωt+ϕ), optimized amplitude (A), angular frequency (ω), and phase shift (ϕ) using the SciPy^58^ library’s curve_fit function. Instances of curve_fit non-convergence were flagged as non-periodical. Additionally, we visually inspected fits for biological relevance, especially for communities with solar year-aligned periods. This approach ensured our findings were both statistically sound and ecologically meaningful.

## Data Availability

Sequencing reads, metagenomic assemblies and MAGs created in this project are deposited under the ENA bioproject PRJEB68319. Information, metadata and accession numbers for all the presented samples and sequence runs are presented in Supplementary Data 1. Taxonomic classifications, quality metrics, accessions and relevant statistics are provided for the 2,332 MAGs in Supplementary Data 2. The number of ResFinder gene counts by sample can be found in Supplementary Data 3. Information on community membership for the MAGs can be found in Supplementary Data 4.

## Acknowledgements

We would like to acknowledge the Novo Nordisk Foundation (NNF16OC0021856), Horizon 2020 grant VEO (874735) and PREPARE-TID (101137132) for funding. We would also like to extend our gratitude to the employees of Fővárosi Csatornázási Művek (Budapest, Hungary) for their efforts in conducting the sewage sampling. We thank for partial support by RRF-2.3.1-21-2022-00006. The Authors wish to also thank Mr. Giuseppe Miccichè and Dr. Marco Bani from ACEA SPA ATO2 WWTP, for logistics and technical assistance when sampling at the WWTP in Rome, Italy as well as Massimo Fulco and Elena Billi from Gruppo HERA for logistics and technical assistance when sampling in Bologna, Italy.

