## Supplementary Figure S1-13 & Supplementary Table S1 for "Time-series sewage metagenomics can separate the seasonal, human-derived and environmental microbial communities, holding promise for source-attributed surveillance"

This document contains the Supplementary Information

#### Supplementary Figures

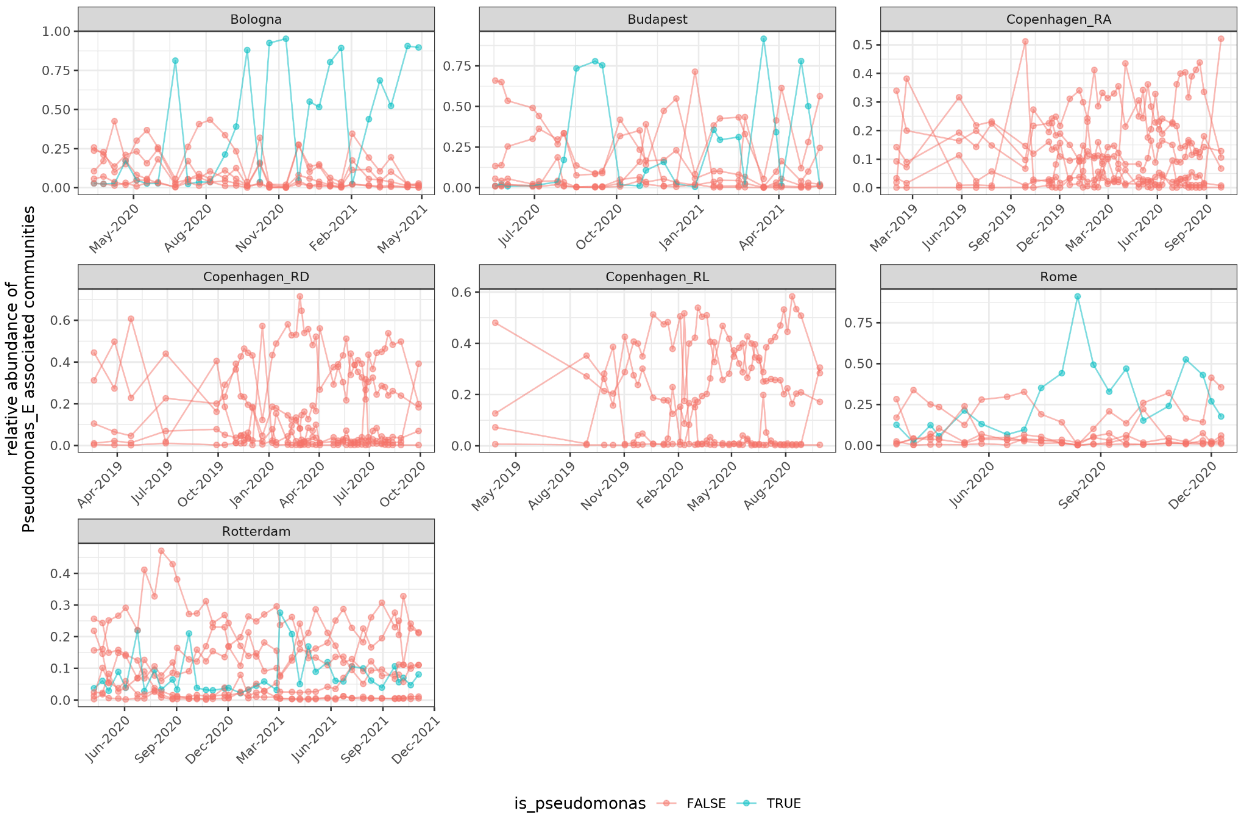


**Figure S1 Timelines of community abundance based on Pseudomonas status.** Sequence depth of MAGs were summed to each community and the CLR transform was computed and shown as a function of whether a community was the most *Pseudomonas_E*-associated.

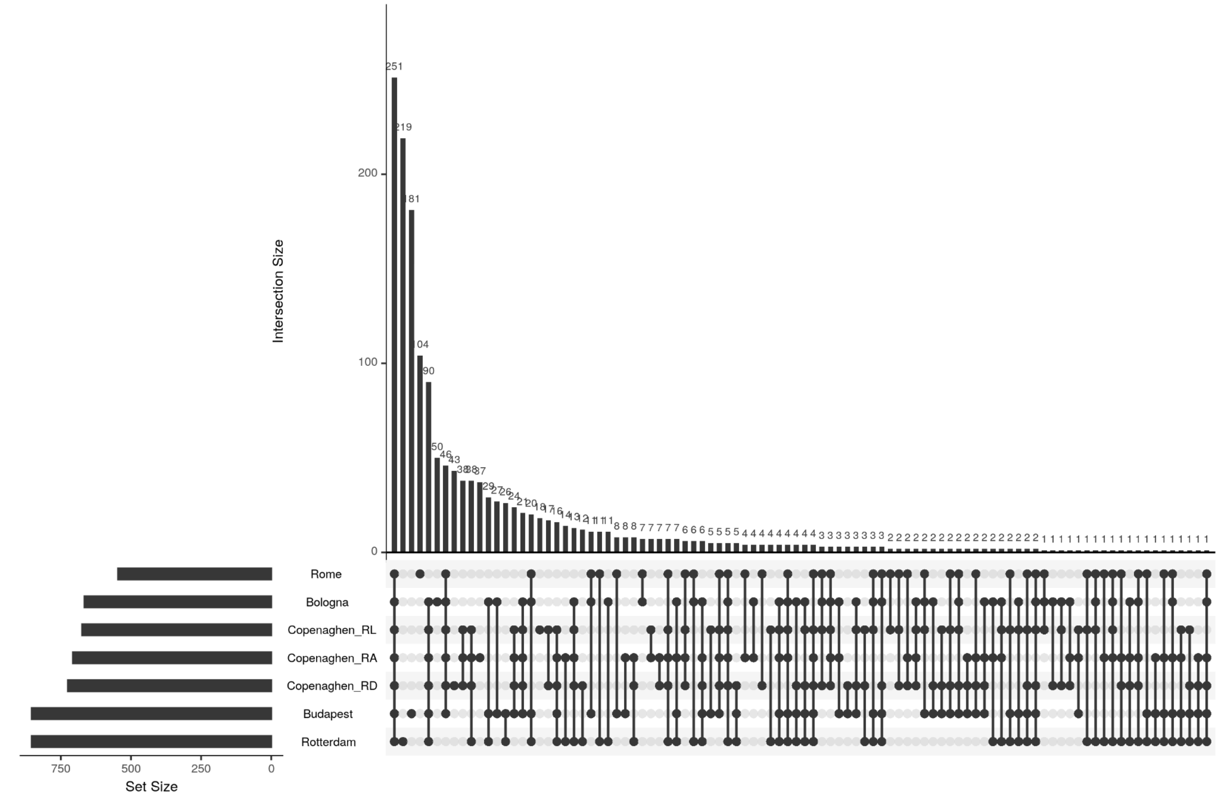


**Figure S2. Preserved Species Across Sewage Sites After the Filtering Step.** This figure, generated using the upsetR package, offers a comprehensive intersectional analysis of microbial taxa preserved across seven sewage sites post-application of a 0.5 mean value threshold. The figure effectively functions as an advanced Venn diagram, clearly depicting shared and unique species across the sites. A notable highlight is the identification of 251 species common across all sites, illustrating a core microbiome component in urban sewage systems. Additionally, site-specific subsets are evident, such as 205 unique taxa in Rotterdam and 180 in Budapest, underscoring the distinctive microbial signatures of each location.

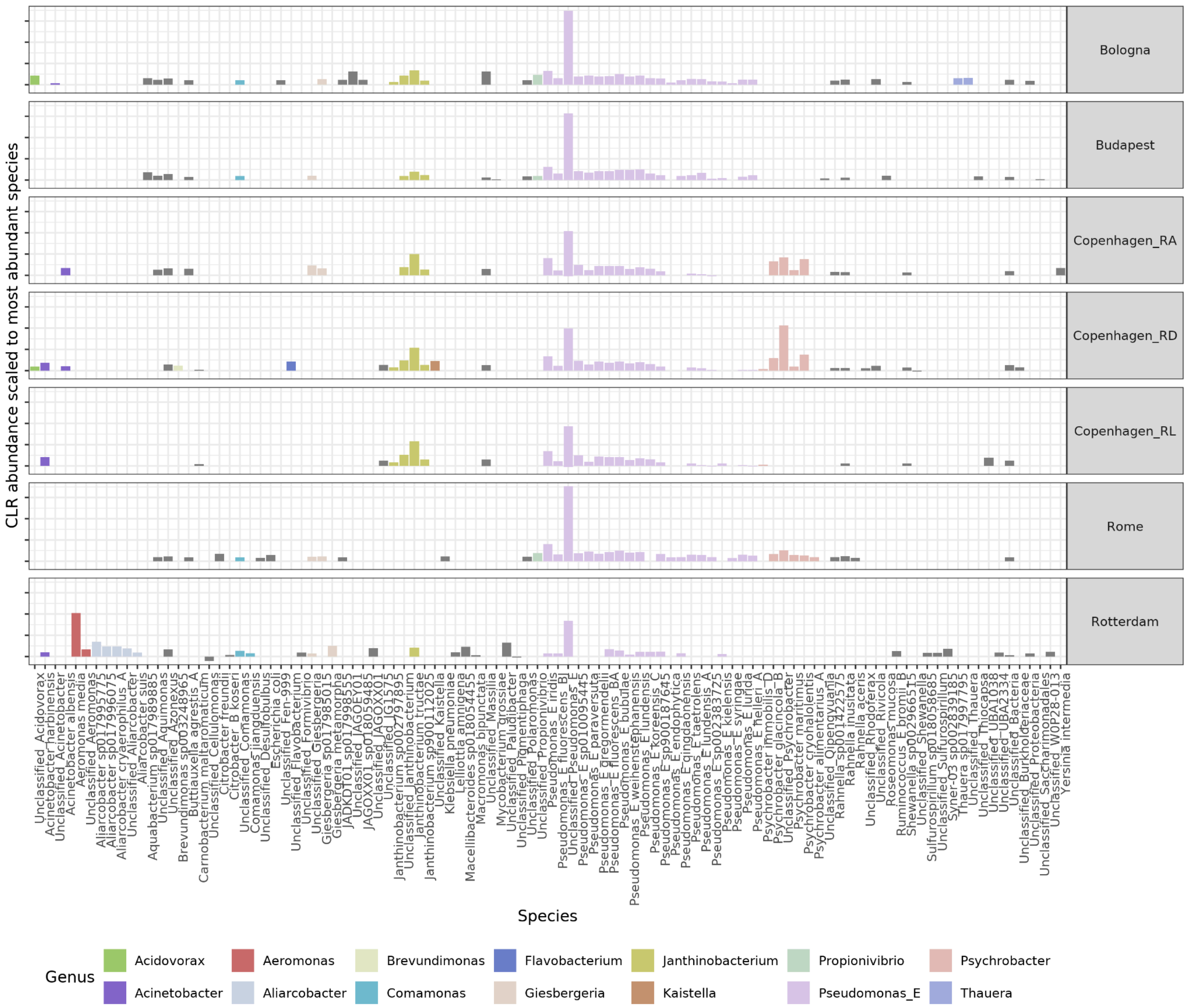


**Figure S3. Differences in species mean CLR abundance between Pseudomonas_E-heavy communities over sites.** *Pseudomonas_E* species-associated community from each sewage site. Within each site, read depths were scaled to the most abundant community member, so other species are a fraction of that. All the non-Rotterdam sites display some shared features but also species-differences.

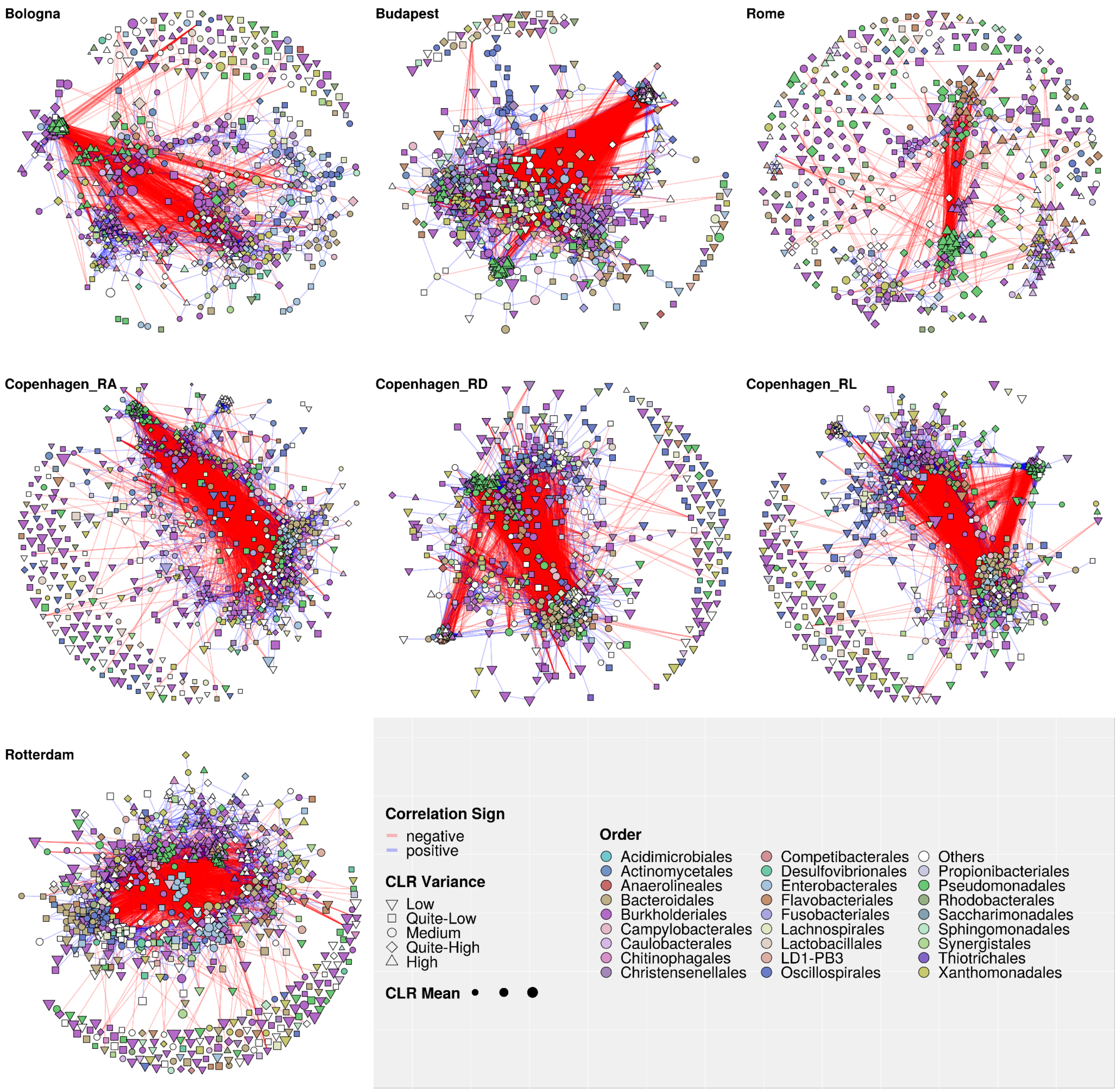


**Figure S4 Taxonomic Order Representation in Microbial Network Diagrams Across Sites:** This figure presents microbial network diagrams for various sewage sites, with each vertex color-coded according to the taxonomic order of the species. Sometimes there is a visible tendency for vertices (species) of the same color (taxonomic order) to cluster together. This pattern suggests a potential facilitation of interactions or co-occurrence among taxonomically similar bacteria. However, it's important to note that this is not a universal rule, as the complex dynamics of microbial communities can lead to interactions beyond taxonomic similarities.

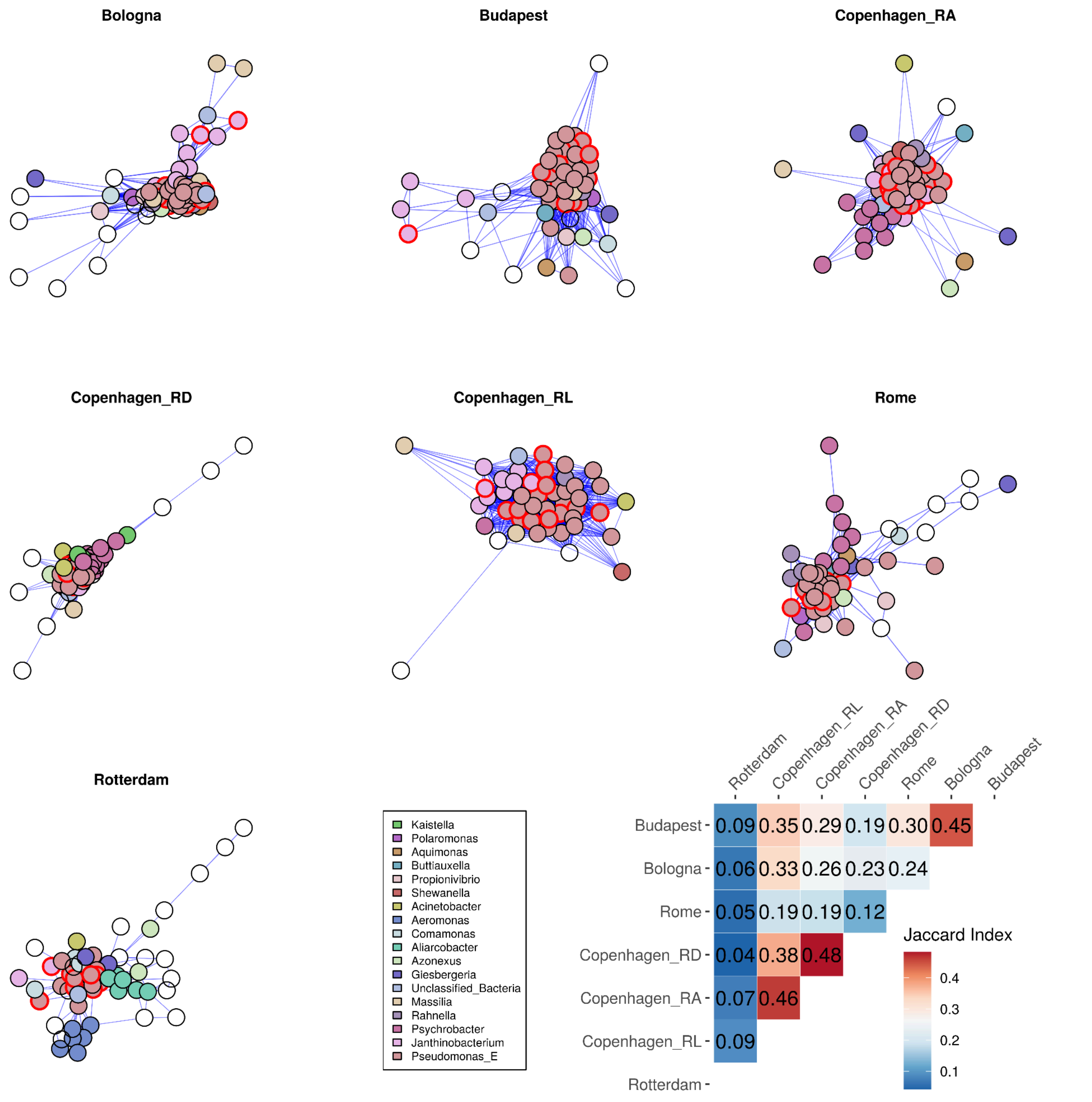


**Figure S5. Network representation of Pseudomonas_E communities across sewage sites.** This figure shows the different *Pseudomonas_E*-dominated communities, represented as subsets of the larger microbial networks. Each network diagram is specific to a site and highlights the Pseudomonas_E species and their interactions. Notably, certain bacterial species within these networks are marked with red borders; these represent the species belonging to the *Pseudomonas_E* genus that are consistently preserved across all the sites after the filtering process.The striking feature of these networks is their high density of connections, suggesting robust and possibly complex interactions within these polymicrobial communities across different urban sewage systems. See **Figure 4** for explanation on the community similarity heatmap.

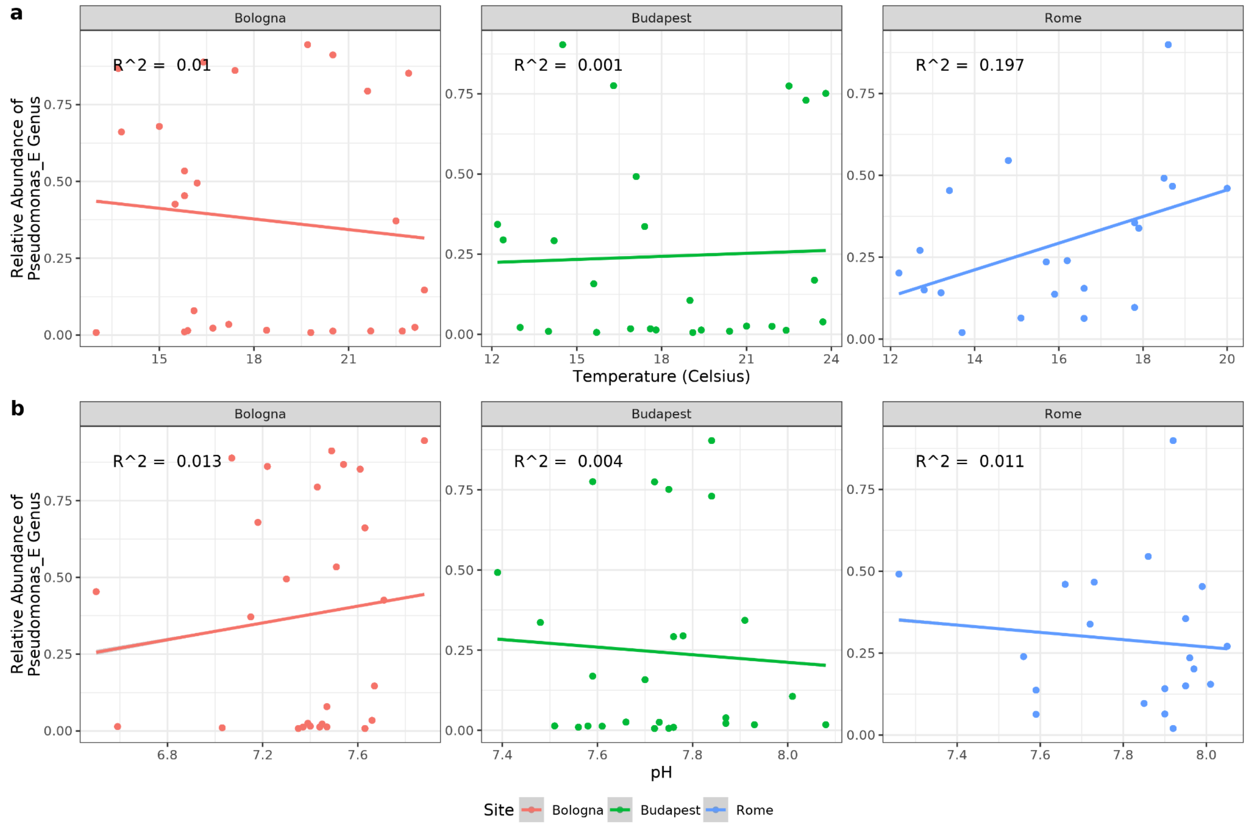


**Figure S6. Comparison of sewage temperature and pH values at sites affected by heavy Pseudomonas_E blooms.** The pH of sewage ranges from 7.5 to 8.0 and remains relatively stable during the sampling period in Budapest and Rome, whereas in Bologna, pH exhibits greater fluctuations. Sewage water temperature varies with the seasons and peaks in the summer. Each panel represents a different sampling site, showing scatterplots of temperature **a**) and pH **b**) versus Pseudomonas_E relative abundance. Linear regression lines and the adjusted R-squared values are shown to each plot, highlighting the general trends observed. We did not detect any striking effects of sewage temperature or pH on the diversity of microbiomes.

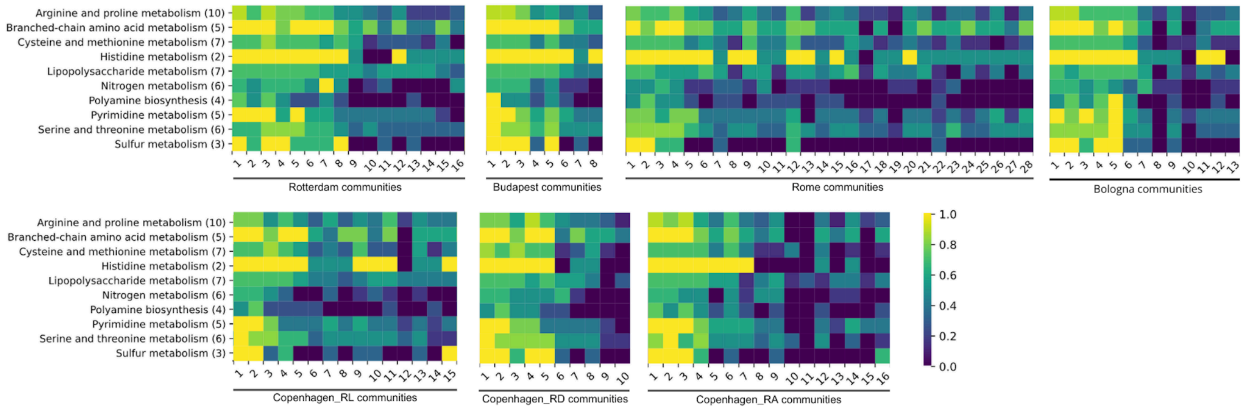

**Figure S7 Metabolic categories with the most variation between distinct communities.** The gradient bar on the right indicates the completeness level of distinct metabolic categories. To the left, the filtered KEGG categories are listed with the corresponding count of KEGG modules contained within each category in parentheses.  As community size increases, different metabolic categories achieve a higher degree of completeness.

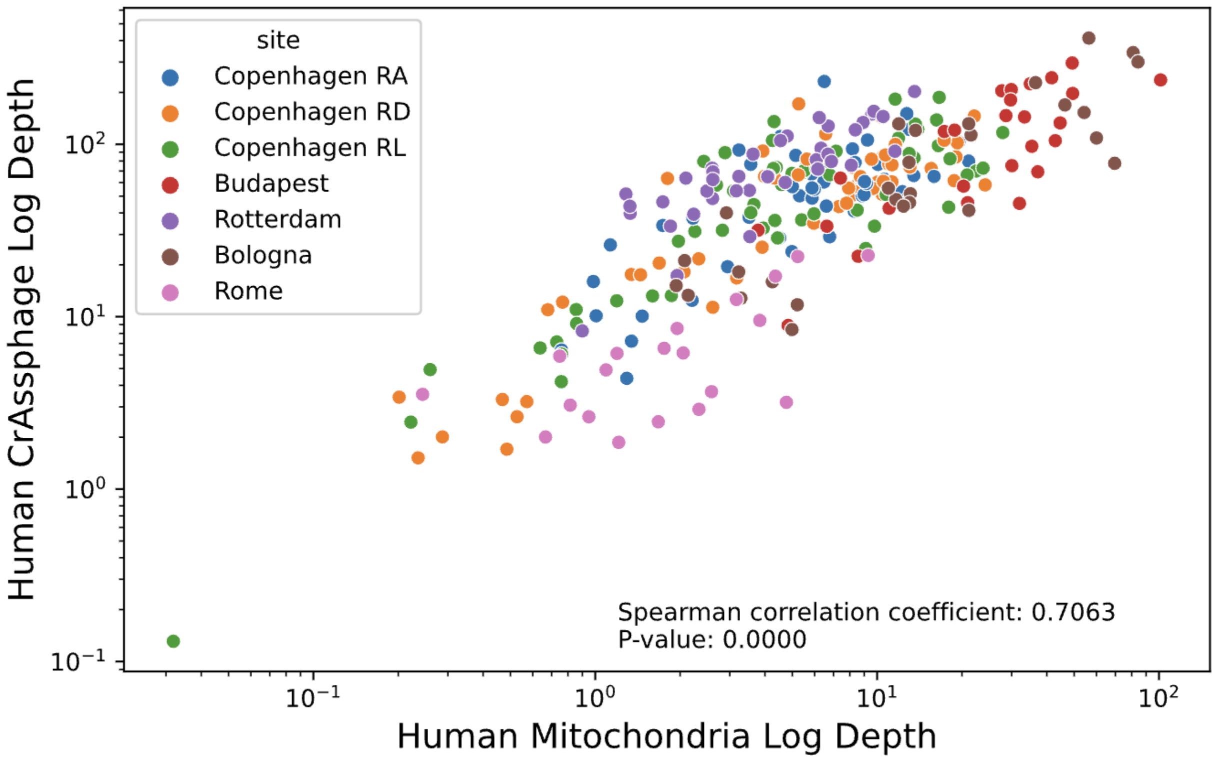


**Figure S8. The presence of the human fecal indicator crAssphage is correlated with the presence of human mitochondria in sewage.** This visualization demonstrates the correlation between the sequencing depths of crAssphage and human mitochondrial DNA.

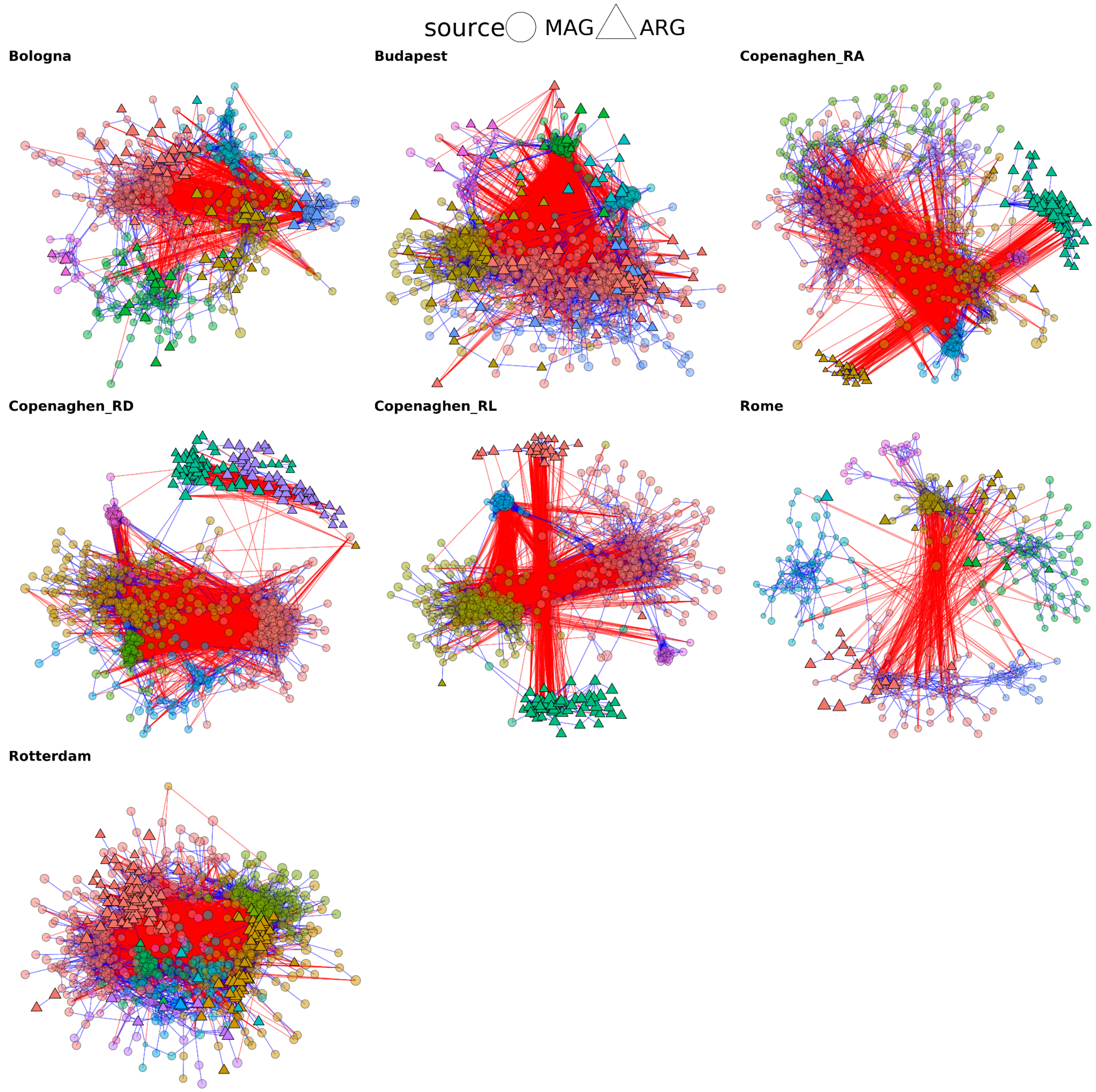


**Figure S9 Community detection in MAG and ARG abundances.** Each vertex corresponds to a species' genome (circle) or ARG (triangle), and the links represent significant positive (blue) and negative (red) correlations. Vertex colors are used to highlight the community membership in combined MAG & ARG networks. For the vertex placement, we used the Fruchterman-Reingold layout algorithm on the subgraph composed from only positive edges. Following this representation, many of the blue/positive links are covered by the circles.

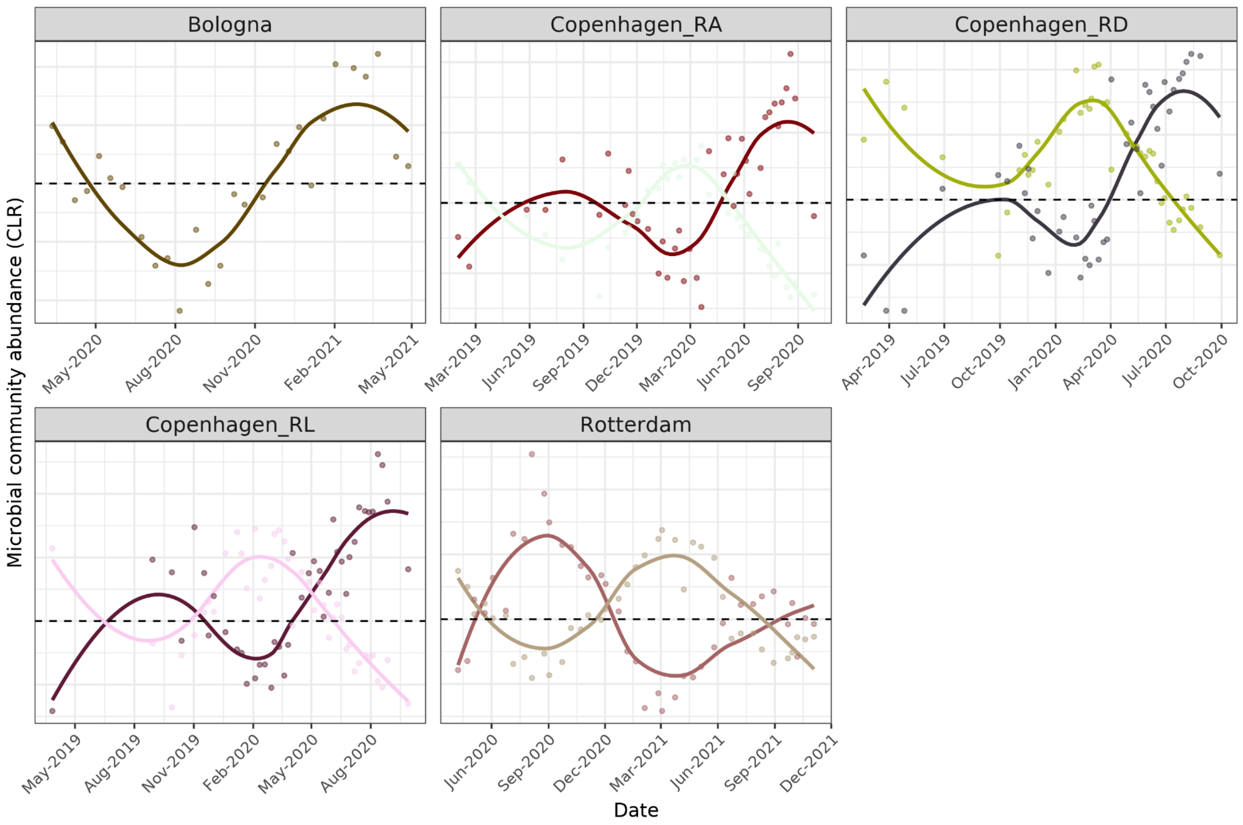


**Figure S10. Annual fluctuations of bacterial communities:** CLR-transformed abundances in bacterial communities exhibiting annual trends, categorized by site. Each scatter plot delineates temporal variations in community abundance, with data points reflecting observed values over time. Overlaying these plots are LOESS lines, highlighting seasonal trends and fluctuations specific to each community. Importantly, the color coding of these plots corresponds to the colors used in the network plots, ensuring a cohesive representation of community dynamics.

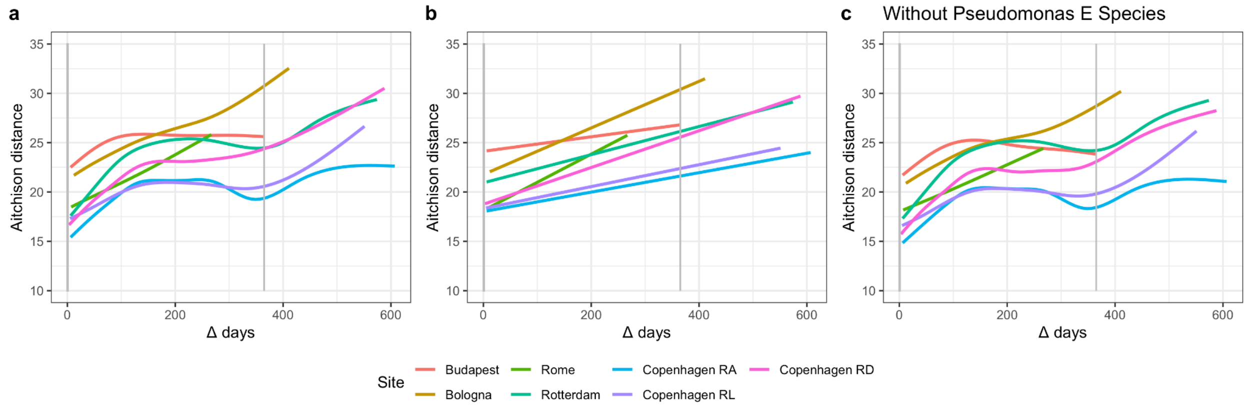


**Figure S11. Sewage microbiomes drift over time and partially return annually.** Within each sampling site, we calculated the beta-diversity (Aichison distance) between any two samples. For each sample pair, the amount of time between sampling (temporal distance) and microbiome distance were computed. Vertical grey lines show whole years in days (0 and 365). **a)** LOESS regression lines show a partial return to similarity for samples taken ~1 year apart. **b)** Linear regression lines showing drift apart (positive slope) for all sites. **c)** Same analysis as '**a**' but the dominant species of *Pseudomonas_E* were omitted prior to the CLR transformation and distance calculation.

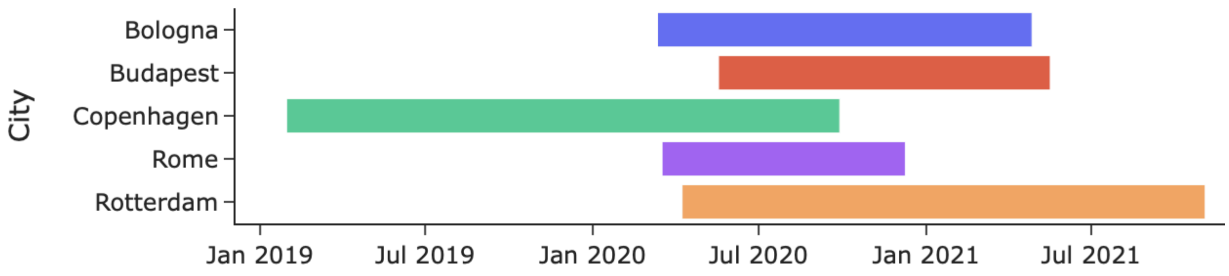


**Figure S12. Sample collection timeline.** Sampling occurred in Bologna from March 12, 2020, to April 27, 2021; in Budapest from May 18, 2020, to May 17, 2021; in Copenhagen from January 30, 2019, to September 28, 2020; in Rome from March 17, 2020, to December 9, 2020; and in Rotterdam from April 8, 2020, to November 3, 2021.

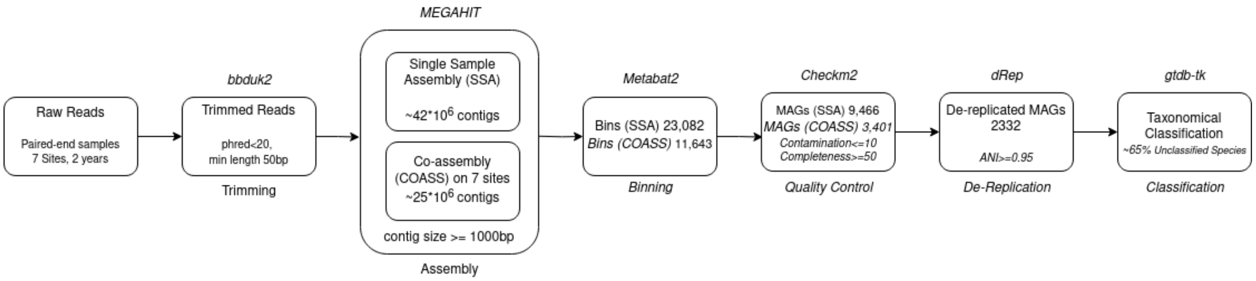


**Figure S13. Workflow of the bioinformatic pipeline for genomic reconstruction, validation and classification.** This diagram provides a concise overview of our genome recovery process, illustrating the tools and the logic employed to achieve our comprehensive dataset.Supplementary Tables

| | Samples | Nodes | Edge Density | Positive Edges | Modularity |
|---|---|---|---|---|---|
| Bologna | 28 | 666 | ~1.99% | ~81% | ~0.60 |
| Budapest | 26 | 854 | ~5.22% | ~70% | ~0.54 |
| CPH RA | 39 | 707 | ~2.48% | ~68% | ~0.54 |
| CPH RD | 42 | 725 | ~4.69% | ~68% | ~0.56 |
| CPH RL | 41 | 674 | ~3.61% | ~69% | ~0.56 |
| Rome | 20 | 546 | ~1.00% | ~81% | ~0.64 |
| Rotterdam | 39 | 854 | ~1.85% | ~75% | ~0.57 |

**Table S1. Summary information on networks of each site.** Information on the specifics of each treatment plant network. The two first columns show the number of included sewage samples and the number of nodes, each represented by a recovered species.

### Supplementary Data

(Please see separate spreadsheet tab)
**Supplementary Data 1.** Table with overview of each of the samples, sequencing runs, metadata and accession numbers in the European Nucleotide Archive.

(Please see separate spreadsheet tab)
**Supplementary Data 2.** Table with overview of the recovered, species-level metagenome-assembled genomes (MAGs) with metadata and accession numbers in the European Nucleotide Archive.

(Please see separate spreadsheet tab)
**Supplementary Data 3.** Table with counts of aligned sequence fragments from the individual metagenomes to the ResFinder database.

(Please see separate spreadsheet tab)
**Supplementary Data 4.** Table with network community information of each MAG in the different sites.

(Please see separate spreadsheet tab)
**Supplementary Data 5.** Table with descriptions for all the columns from the other tables.
